# TXA11114: Discovery of an *in vivo* efficacious efflux pump inhibitor in *Pseudomonas aeruginosa*

**DOI:** 10.1101/2025.02.28.640909

**Authors:** Jesus D. Rosado-Lugo, Pratik Datta, Ahmad Altiti, Yongzheng Zhang, Jun Lu, Yi Yuan, Ajit K. Parhi

## Abstract

Multi-drug resistance in *Pseudomonas aeruginosa* is often associated with overexpression of drug efflux pumps which limit antibiotics exposure. So far, successful development of efflux pump inhibitors (EPIs) has been plagued by undesirable toxicities and inconsequential *in vivo* efficacy. TAXIS Pharmaceuticals Inc. has discovered an effective anti-pseudomonal therapy involving a novel indole carboxamide class of EPI, TXA11114, with a fluorine substituted diamine sidechain, as an adjunctive to levofloxacin. TXA11114 has demonstrated excellent potentiation of levofloxacin MIC by ≥ 8-fold in 90% of Walter-Reed and CDC multi-drug resistant (MDR) isolates. Biophysical and genetic studies with TXA11114 support efflux inhibition while ruling out membrane disruption as a mechanism of action. TXA11114 enhanced the levofloxacin killing and diminished the frequency of resistance emergence to levofloxacin to undetectable levels. Moreover, in murine thigh and lung *P. aeruginosa* infection models, the TXA11114-levofloxacin combination showed pronounced killing compared to levofloxacin alone, achieving a validated *in vivo* efficacy milestone that previous EPIs could not. Most importantly, TXA11114 exhibits a safe toxicology profile when screened for cytotoxicity, hERG channel inhibition, *in vitro* nephrotoxicity, and acute toxicity. Further, pharmacokinetic (PK) parameters of TXA11114 have a complementary profile with that of levofloxacin in plasma and bronchoalveolar lavage fluid (BALF) samples of infected mice, maximizing pharmacodynamic (PD) benefits. Overall, studies on the TXA11114-levofloxacin combination highlight its potential as an anti-pseudomonal agent for combating multidrug-resistant infections.

## Introduction

Centers for Disease Control and Prevention (CDC) and the World Health Organization (WHO) have declared antimicrobial resistance (AMR) to be one of the most serious problems facing our national and global health systems today. As per a CDC 2019 report, AMR is attributable to 1.27 million killing worldwide and associated with 5 million deaths. In the US alone, each year 2.8 million AMR infections cause 35,000 mortalities [1]. A recent CDC report from 2024 highlights a further increase in antimicrobial resistance in the post-COVID-19 pandemic period compared to pre-pandemic levels [2]. Gram-negative bacteria are generally more resistant to antibiotics due to their unique cell envelope structure. Additionally, the presence of multiple resistance mechanisms drives multidrug resistance, frequently resulting in treatment failures in clinical settings [3].

*P. aeruginosa*, an opportunistic Gram-negative pathogen, is known for its adaptability and is considered as one of the paradigms of antimicrobial resistance. It is a leading cause of hospital-acquired and chronic infections, contributing to significant morbidity and mortality and posing a serious public health threat [4]. In 2015, the incidence rate of *P. aeruginosa* was 32.6 per 100,000 persons per year in the Military Health System (MHS). This rate reflects a 13.6% increase from the weighted historic baseline in the preceding three years. The South and West MHS regions had the highest incidence rates (43.0 per 100,000 persons per year and 39.9 per 100,000 persons per year, respectively), but the South and South Atlantic MHS regions had the highest incidence of multidrug-resistant (MDR) infections (3.8 per 100,000 persons per year and 2.2 per 100,000 persons per year, respectively). Among all MHS beneficiaries, 47.0% of *P. aeruginosa* infections were healthcare-associated cases. *P. aeruginosa* infections did not display 100.0% susceptibility to any tested antibiotic in the MHS in 2015. The most prescribed antibiotic classes associated with prevalent *P. aeruginosa* infections in 2017 were fluoroquinolones (55.1%), penicillins and inhibitors (16.6%), and cephalosporins (10.7%). Within the fluoroquinolone class, ciprofloxacin (33.5%) and levofloxacin (21.6%) were the most prescribed ones, accounted for more than 50% of all prescriptions associated with *P. aeruginosa* infections in the MHS, and both had an efficacy of less than 90% [5].

Antimicrobial resistance in Gram-negatives is governed by four main mechanisms, including limiting the uptake of a drug, altering a drug target, inactivating a drug, and active drug efflux [3]. Overexpression of multidrug drug efflux pumps can expel several unrelated classes of antibiotics, promoting intrinsic and acquired resistance in *P. aeruginosa* [6–9]. In Gram-negative bacteria, many of these efflux pumps belong to the resistance-nodulation-cell division (RND) family of tripartite efflux pumps [10, 11]. The RND efflux systems in *P. aeruginosa* consist of many identified efflux pumps that are largely responsible for its multidrug resistance [7]. Four well characterized multidrug efflux pump systems (MexAB-OprM, MexCD-OprJ, MexEF-OprN, and MexXY-OprM) are prevalent in clinical isolates of *P. aeruginosa*. Overexpression of *mexB*, *mexF* and *mexY* was detected in 27, 12, and 45% of 33 *P. aeruginosa* clinical isolates tested, respectively [9]. Additionally, *P. aeruginosa* possesses six RND efflux pumps (MexJK, MexGHI-OpmD, MexVW, MexPQ-OpmE, MexMN, and TriABC-OpmH) which might contribute to resistance at the clinic [12–15]. Approaches for circumventing efflux-mediated resistance can be direct or indirect. For example, a direct approach is to develop new antibiotics that are poor substrates of efflux pumps. An indirect approach involves developing efflux pump inhibitors (EPIs) that need to be paired with antimicrobials that have become ineffective by efflux. Due to dwindling pipelines of new antibiotics, the inhibition of efflux pumps has become an attractive avenue to rejuvenate older antimicrobials to tackle the antibiotic resistance problem [15, 16]. An ideal EPI would efficiently block these pumps, increasing the concentrations of the antimicrobials within the cell, thus rendering them effective again [17]. Identification of EPIs with *in vivo* efficacy against *P. aeruginosa* holds promise for developing a successful combination therapy for MDR infections, which has so far remained unsuccessful in the clinic. Reducing the minimal inhibitory concentration (MIC) of an established anti-pseudomonal agent such as levofloxacin in resistant isolates to below the clinical breakpoint should make it possible to use this drug routinely again. Moreover, coadministration of an EPI will lower the MIC for levofloxacin in susceptible isolates that are currently treatable with levofloxacin alone and reduce the likelihood that rare resistant mutants will emerge. Therefore, inhibition of efflux pumps may help solve the antibiotic resistance problem in multiple ways, by restoring the efficacy of older antimicrobials and by contributing to reducing the emergence of resistance. Lastly, the first-in-class use of EPIs obviates the need to discover new antibiotics, a strategy that saves a lot of time, effort, and capital associated with the discovery and development of new antibiotics.

A variety of chemical scaffolds have been shown to act as EPIs (Figure 1) [18–21]. Among the first EPIs is the peptidomimetic MC-207,110 (Phenylalanine-arginine β-naphthylamide, PAβN), and related dipeptide amide compound, MC-04,124. Both were developed by Essential Therapeutics Inc. and have shown to potentiate levofloxacin effectively in wild-type and efflux-pump-overexpressed *P. aeruginosa* strains [13, 14]. A non-peptidomimetic lead series worthy of mentioning is D13-9001, the lead candidate from a novel pyridopyrimidine scaffold-based series developed for MexAB-OprM specific RND-pump inhibition [15]. A more recent endeavor involves MBX-2319, from a pyranopyridine class of compounds that potentiates ciprofloxacin, levofloxacin, and piperacillin against *E. coli* [16]. However, the development of these compounds has stalled or even stopped for various reasons. PAβN and related compounds showed prolonged accumulation in tissues associated with renal toxicity [17]. MC-04,124 was discontinued following the closure of Essential Therapeutics Inc. [18]. The MBX EPI series does not inhibit efflux in *P. aeruginosa* and, thus, may not be suitable for anti-pseudomonal therapy [19]. Despite exhibiting good *in vitro* activity and being efficacious *in vivo* against *P. aeruginosa*, D13-9001 is yet to progress to clinical evaluation [20]. Thus, the discovery of novel EPIs active against Gram-negative pathogens, particularly *P. aeruginosa*, with the potential of clinical use is still needed.

**Figure 1.**
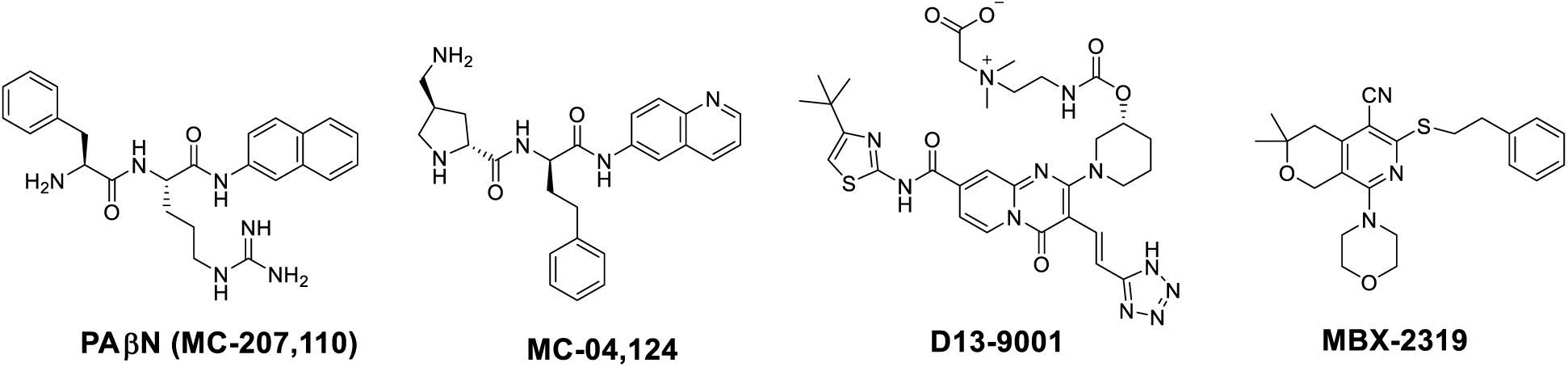
Previously known efflux pump inhibitors

TAXIS Pharmaceuticals Inc. is working to find solutions to efflux mediated multidrug-resistance by developing safe and efficacious EPIs that truly hold potential as a first-in-class treatment option [22]. We have previously published preliminary efforts on a diaminopentanamide class of potentiators and more recently on heterocyclic carboxamide classes of potentiators that yielded TXA01182 [23] and its constrained analog TXA09155 (Figure 2) [24]. Both these novel EPIs potentiate monobactam, fluoroquinolones, sulfonamides, and tetracyclines against *P. aeruginosa* at a minimum concentration of 6.25 μg/mL. Further, TXA01182 and TXA09155 demonstrated a synergistic effect with levofloxacin against several MDR *P. aeruginosa* clinical isolates, the latter being more potent than TXA01182. Both EPIs possess two amine groups, which are crucial for their activity. Removing both or one of them completely alienates their potentiation effect. Whilst basic centers are attractive features to have in drug molecules for a variety of reasons including aqueous solubility and interactions with binding sites, they often are accompanied by problems including off-target liabilities such as hERG, CYP inhibition and reduced permeability [25]. Tissues containing cells rich in lysosome vacuoles are particularly sensitive to the toxic effects of intralysosomal accumulation of basic molecules [26]. Nephrotoxicity due to intralysosomal accumulation in renal tissue is a widely documented phenomenon [27]. This may be the major reason why PAβN and related series of compounds did not progress beyond the preclinical stage. The basic amine functionalities in the PAβN class of EPIs were found to be associated with unfavorable pharmacokinetic and toxicological profiles, and development of this lead series was suspended back in mid-2000s. We are aware of these potential liabilities in our pharmacophores which require two amine groups to exert their activity. The aim of this study is to investigate the effect of reducing the basicity of the amines and its impact on the potentiation activities and the toxicity profile of this novel EPI series.

**Figure 2.**
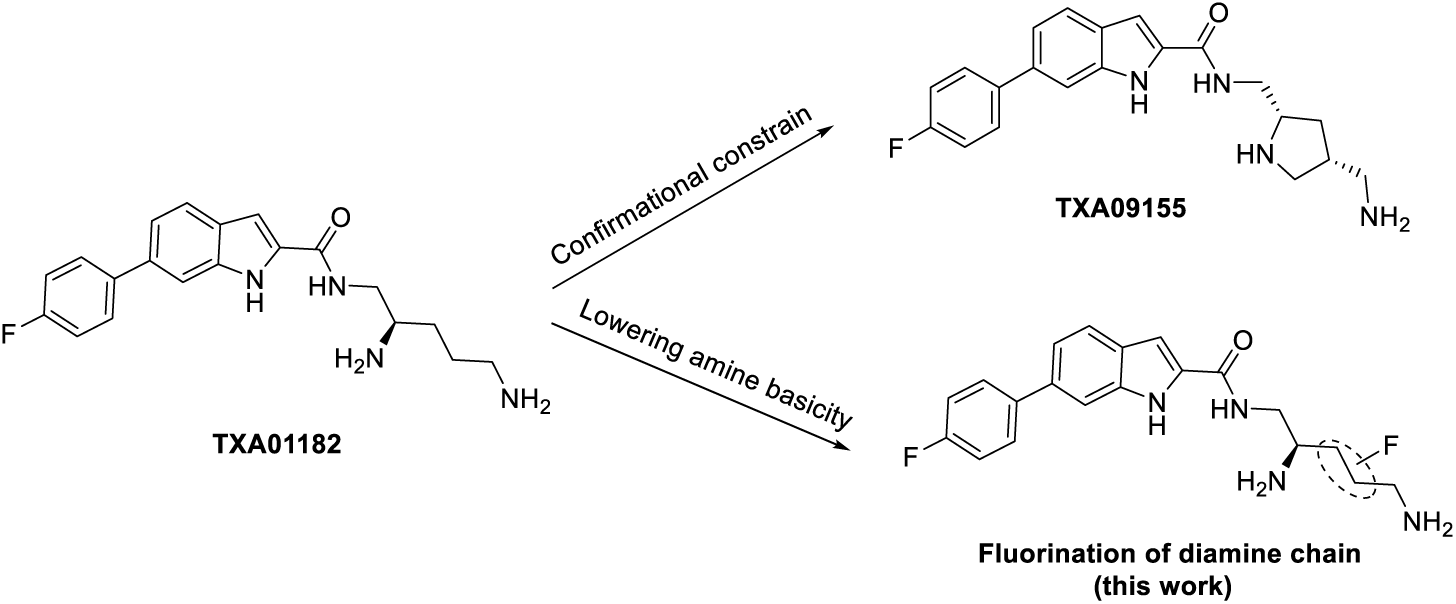
Sequential progression of TAXIS’ carboxamide EPIs

Strong electron-withdrawing groups such as nitriles, esters, amides, ketones, oxetanes, and polyheteroatom heterocycles can modulate the pKa of amines but may introduce additional pharmacophoric elements. Alternatively, replacing hydrogen with fluorine in drug molecules can enhance their properties. Due to fluorine’s similar size to hydrogen, such substitutions typically have minimal structural impact. However, fluorine’s distinct electronic properties can create new interactions with the drug’s target, potentially enhancing activity.

Fluorine has been widely used in drug design and development for its ability to influence pKa, enhance intrinsic potency, and improve metabolic and pharmacokinetic (PK) properties. In our pharmacophore side chain, the pKa of both amines can be adjusted through the strategic placement of fluorine atoms between them. Mono- and di-substituted fluorine modifications at two possible carbon positions between the amines can alter their pKa depending on the number and location of substitutions.

This study focuses on reducing the basicity of the two amines in our EPI molecules by selectively fluorinating the diamine side chain and evaluating the impact of these modifications on EPI activity and toxicity. Accordingly, fluorination of the amine side chains in TXA01182 was performed, yielding three fluorine-substituted EPIs (Figure 3).

**Figure 3.**
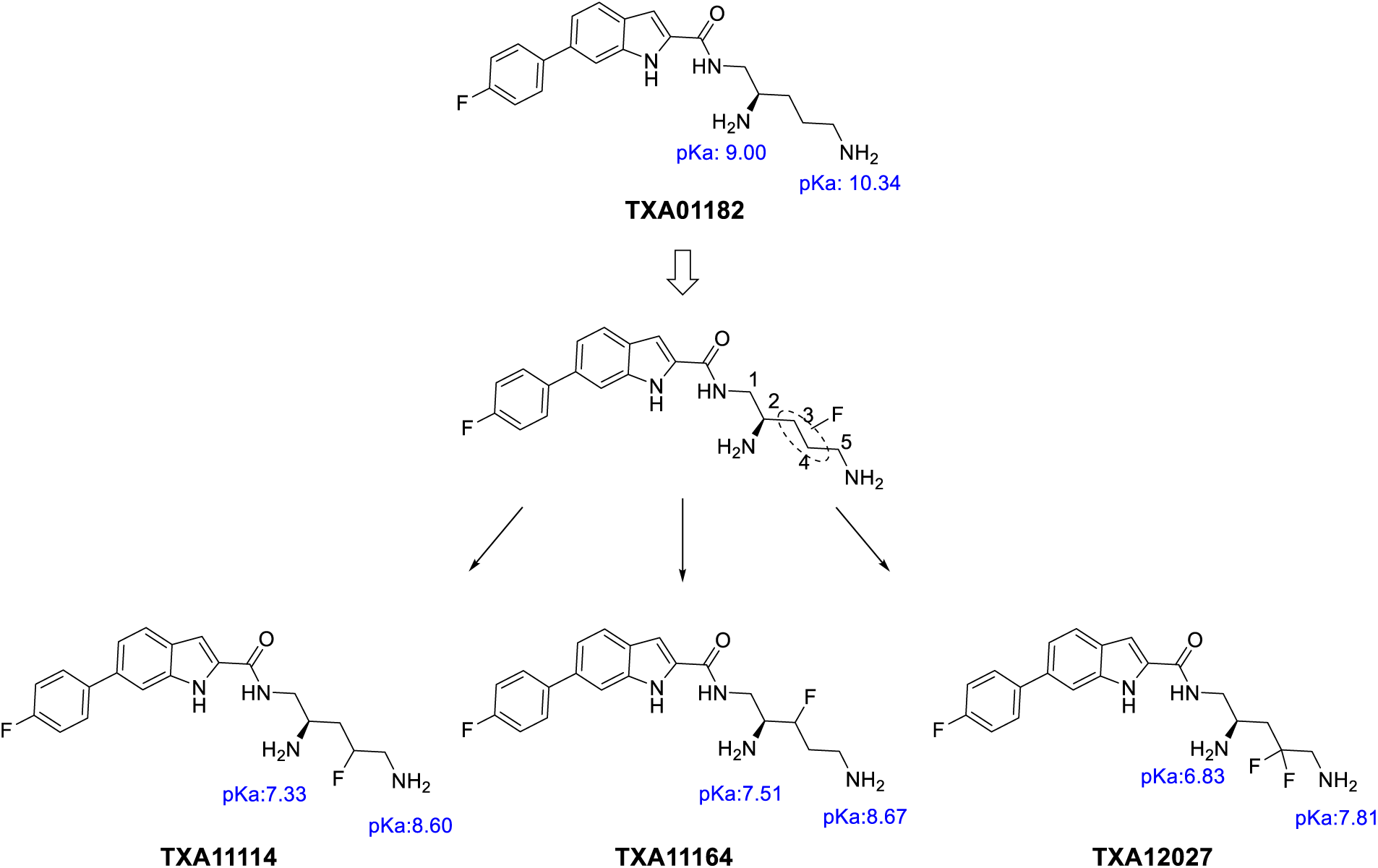
Structures of three TAXIS EPIs containing fluorinated side chains.

The mono-fluorination at C-3, C-4, or di-fluorination at C-4 of the diamine side chain yielded three unique EPI molecules: TXA11114, TXA11164 and TXA12027 which were deemed sufficient to investigate the influence of the fluorine substitutions on the diamine side chain of these EPIs. The pKa of amines (calculated from ACD/LC simulator) owing to the F-substitutions at either C-3 or C-4 are shown. Di-fluorination at C4 in TXA12027 resulted in a pKa reduction greater than 2 units compared to TXA01182. Mono-fluorination at C-3 or C-4 reduced the pKa by approximately 1.5 units in TXA11114 and TXA11164, respectively, compared to TXA01182.

## Results and Discussion

### 1. Synthesis of fluorinated EPI analogs

To study the fluorine-induced modulation of the pKa of the amine groups in the proposed EPIs, our syntheses feature the selectively introducing fluorine moieties vicinal to each amine group. Therefore, we invested in the accessible intermediates **2**, **3**, and **4** (scheme1), all are made from commercially available *S*-Garner aldehyde **1**. Applying the HWE reaction on aldehyde **1** according to literature [31] using triethyl 2-fluoro-2-phosphoacetate and LiHMDS afforded **2** as an E/Z mixture, hydrogenation of the double bond in **2** using Pt/C followed by a sequence of NaBH_4_ reduction of the ester moiety, Mitsunobu amination of the resulting alcohol, and finally, protecting group manipulation afforded alcohol **5** in 39% over 4 steps. The allylation of Garner’s aldehyde **1** using allyl bromide and activated Zn dust at 65 °C affords the corresponding allyl alcohol as a mixture of diastereomers [28], which was converted to allyl fluoride **3** at −78 °C by nucleophilic fluorination using DAST [30] in 62%. Compound **3** was found as a mixture of diastereomers epimeric at the fluorinated center; a cleavage of the acid labile protecting groups in a small sample of **3** revealed that the diastereomeric ratio of the fluorinated derivatives is 4:1 based on the ^1^HNMR analysis (*Supporting information*). Oxidative cleavage of the double bond in **3** with NaIO_4_/K_2_OsO_4_.H_2_O, followed by NaBH_4_ reduction, Mitsunobu amination, and protecting group manipulations afforded the fluorinated diamine alcohol **6** in 68 % yield with fluorine atom installed vicinal to the proximal amine group at C3. A subsequent Mitsunobu amination followed by hydrazine cleavage of the corresponding phthalimide furnished the triamine derivatives **8** and **9** in 75% and 53% yield over two steps, respectively. The difluoro ester derivative **4** [66, 67] was reduced by NaBH_4_ in the THF/MeOH to give the corresponding alcohol; functionalizing the resulting alcohol using the Mitsunobu reaction seemed ineffective. Alternatively, the alcohol was converted to the corresponding triflate, which was subsequently subjected to Gabriel amine synthesis using sodium phthalimide at 125 ^0^C, which afforded the desired compound in a 53% yield over two steps. Functional group manipulations on the resulting phthalimide provided an intermediate alcohol **7** in 27% yield over five steps. The same amination protocol used in the previous two examples was applied to intermediate **7** to accomplish the amine **10** in 80% with difluoride at C4, vicinal to the distal amine moiety.

**Scheme 1.**
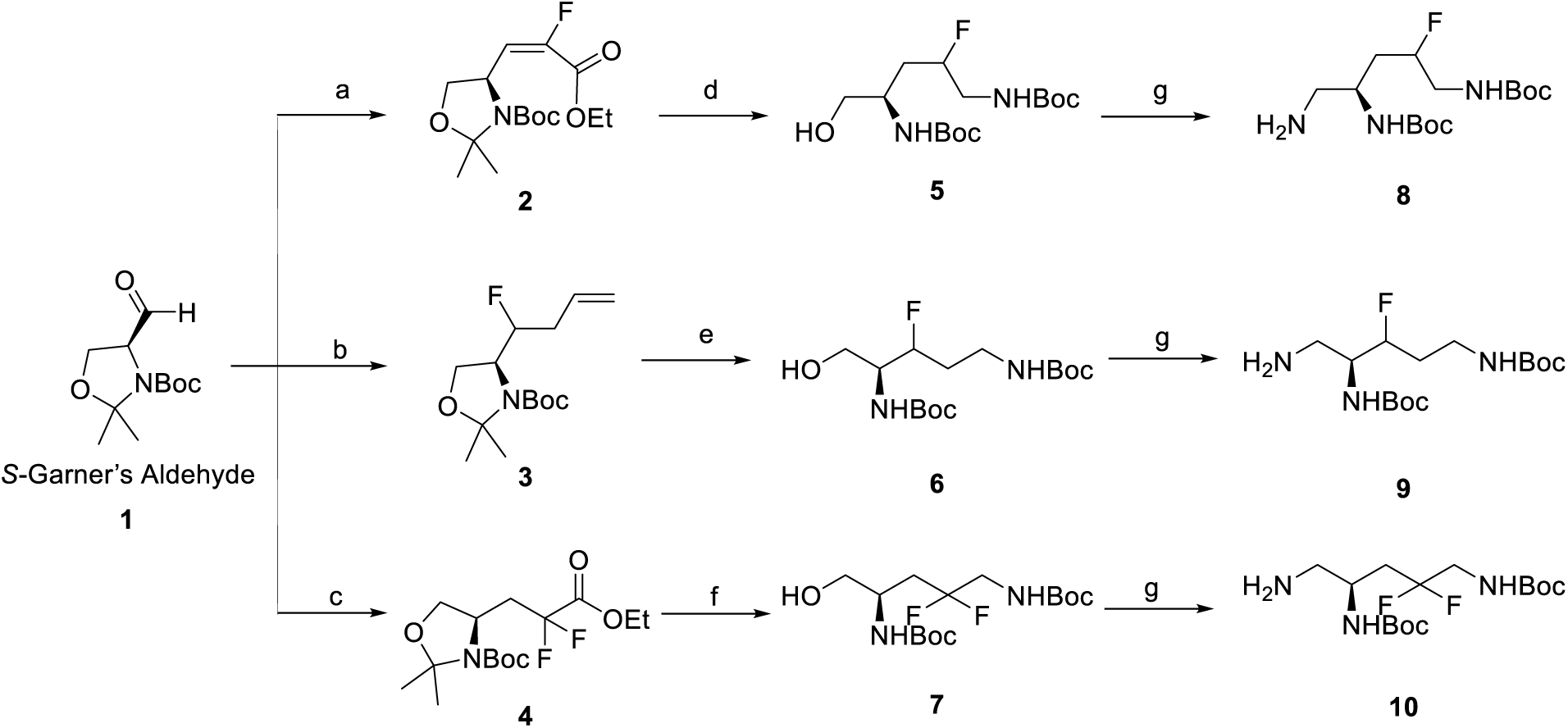
Reagents and reaction conditions: a) Reference [31]: Triethyl 2-fluoro-2-phosphoacetate LiHMDS, THF, −40 °C -rt, 16h, (78%). b) Reference [28]: 1. Allyl bromide, Activated Zn dust, THF 50 °C, 6h (65%); 2. DAST, DCM, −78 °C - rt, 16 h (64%). c) Reference [66, 67]: 1. BrCF_2_CO_2_Et, Zn, ultrasounds, THF, rt, 8 h.; 2. TCDI, DCE, rt, 20 h; 3. Et_3_SiH, Bz_2_O_2_, reflux 1.5h. d) 1. H_2_ gas (1 atm), Pt/C, MeOH; 2. LiBH_4_, MeOH; 3. Phthalimide, Ph_3_P, DIAD, THF, 0 °C-rt (2h); 4. NH_2_NH_2_.H_2_O, EtOH, 90 °C (2h); 5. 4 M HCl/ dioxane, MeOH, 50 °C, 2h; 6. Boc_2_O, Et_3_N, DCM, 4h. e) 1. K_2_OsO_4_, NaIO_4_, THF-H_2_O, 16h, 0 °C-rt; 2. NaBH_4_, MeOH, rt (20 min); 3. Phthalimide, Ph_3_P, DIAD, THF, 0 °C-rt (2h); 4. NH_2_NH_2_.H_2_O, EtOH, 90 °C (2h); 5. 4 M HCl (dioxane), MeOH, 50 °C, 2h; 6. Boc_2_O, Et_3_N, DCM, 4h (68% over 7 steps). f) 1. NaBH_4_, THF-MeOH (5:1), 0 °C -rt (2h); 2. Tf_2_O, Pyridine, DCM, 0 °C -rt 45 min, then Phthalimide, NaH, DMF, 125 °C, 24 h; 3. NH_2_NH_2_.H_2_O, EtOH, 90 °C (2h); 4. 4 M HCl (dioxane), MeOH, 50 °C, 2h; 5. Boc_2_O, Et_3_N, DCM, 4h. g) 1. Phthalimide, Ph_3_P, DIAD, THF, 0 °C-rt (2h); 2. NH_2_NH_2_.H_2_O, EtOH, 90 °C (2h).

Having successfully the desired fluorinated amines intermediates in hand, we turned our attention to finishing the synthesis of new EPIs. Accordingly, TXA1164 and TXA12027 were made in 82% and 85%, respectively using HATU coupling followed by Boc groups deprotections using TFA. TXA11114 was made in 78% using EDC/ HOBt as coupling reagents followed by Boc deprotection using 4 M HCl solution.

**Scheme 2.**
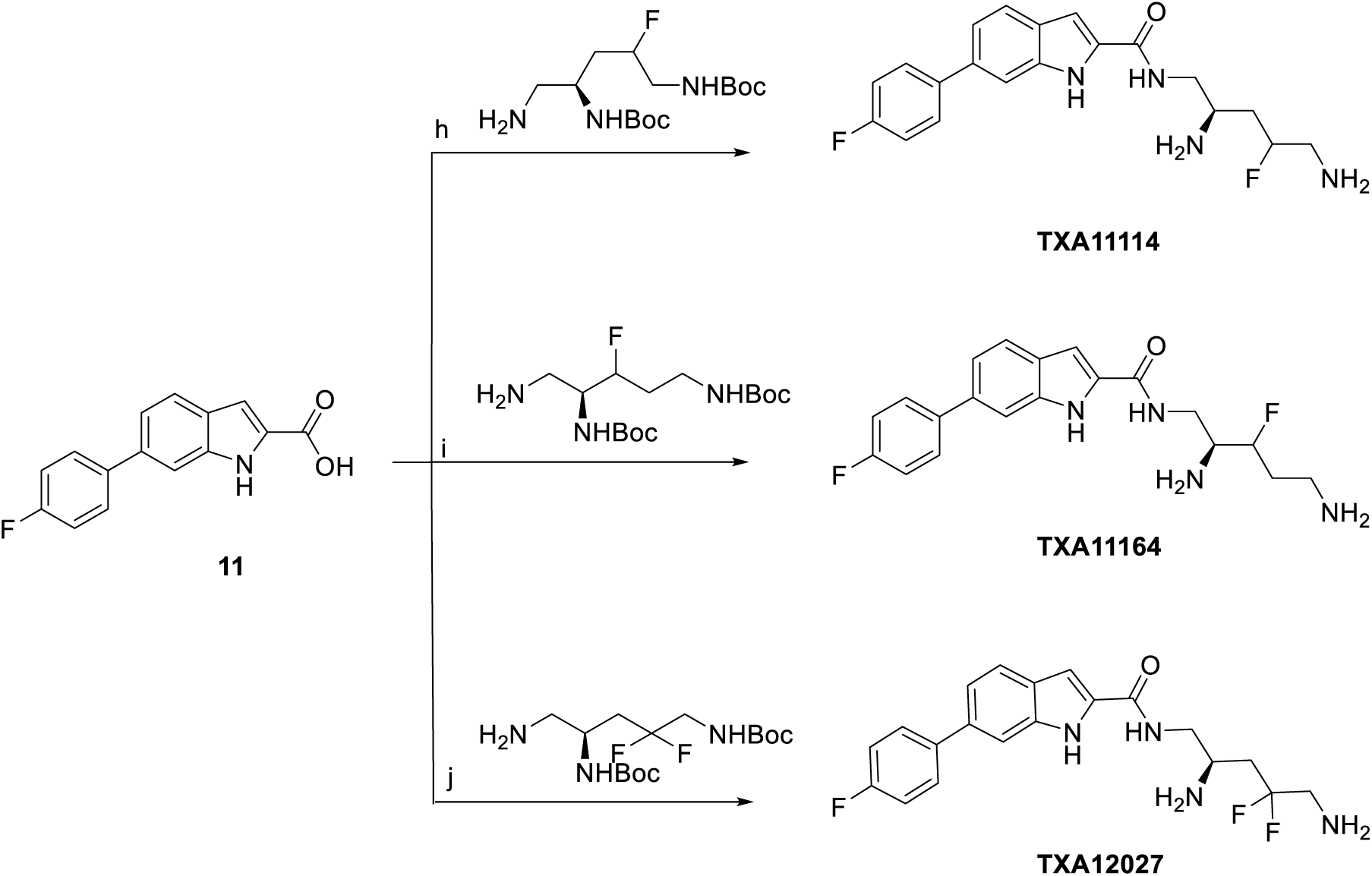
Reagents and conditions: h) 1. Amine **8**, EDC, HOBt, DMF. 2. 4M HCl (dioxane) (78%). i) 1. Amine **9**, HATU, DMF, Et_3_N, 20 min, 2. 95%TFA, 2.5% Et_3_SiH, 2.5% H_2_O. (82%), j) 1. Amine **10**, HATU, DMF, Et_3_N, 20 min. 2. 95%TFA, 2.5% Et_3_SiH, 2.5% H_2_O (85%).

### 2. Evaluation of the fluorinated EPIs for their potentiation of levofloxacin

Once synthesized, the three fluorinated EPI analogs (TXA11114, TXA11164 and TXA12027) were first assayed for their antibacterial activities against *P. aeruginosa* ATCC 27853. They had MICs of 100, >200 and 100 µg/mL, respectively (Table 1). The potentiation abilities of these three fluorinated compounds with levofloxacin were then determined at 6.25 μg/mL and compared with two first generation EPIs: TXA01182 and TXA09155 [23, 24]. Among the three EPIs, only TXA11114 could potentiate levofloxacin by 8-fold comparable to TXA01182 and TXA09155 (Table 1). Strikingly, TXA11164 and TXA12027 failed to potentiate levofloxacin. Particularly, inactivity of TXA11164 was surprising as the pKa values of both the amines are similar to TXA11114. Based on our previous findings, EPIs that are inactive in combination with levofloxacin have often remained inactive in other combinations also, prompting the exclusion of TXA11164 and TXA12027 from further analysis [23, 24].

**Table 1.**
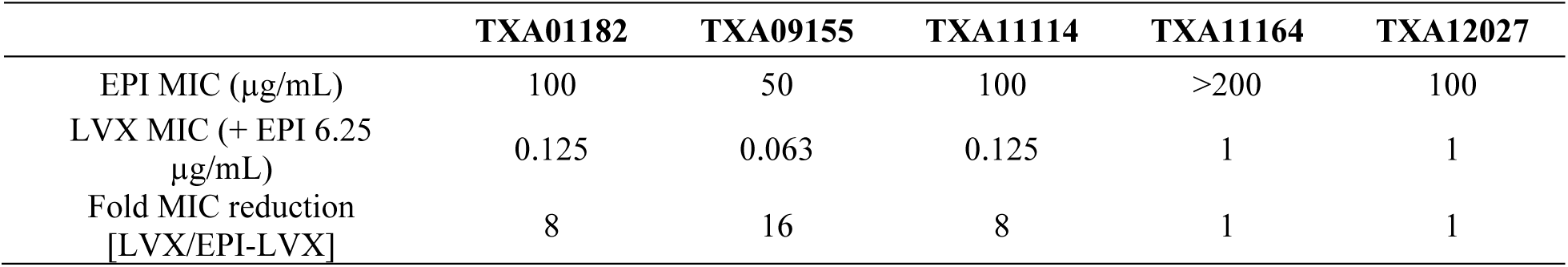
Comparison of potentiation abilities of fluorinated EPI analogs in combination with levofloxacin against *P. aeruginosa* ATCC 27853. LVX: levofloxacin

|  | TXA01182 | TXA09155 | TXA11114 | TXA11164 | TXA12027 |
| --- | --- | --- | --- | --- | --- |
| EPI MIC (µg/mL) | 100 | 50 | 100 | >200 | 100 |
| LVX MIC (+ EPI 6.25 µg/mL) | 0.125 | 0.063 | 0.125 | 1 | 1 |
| Fold MIC reduction [LVX/EPI-LVX] | 8 | 16 | 8 | 1 | 1 |

LVX: levofloxacin

### 3. TXA11114 potentiates other antibiotics with efflux liabilities

TXA11114 potentiation of levofloxacin warranted its further evaluation in combination with other antibiotics, as shown in Table 2. The ability of TXA11114 to lower the MIC of multiple antibiotics with efflux liabilities in *P. aeruginosa* was tested at concentrations ranging from 25 to 3.13 µg/mL (1/4^th^ to 1/32^nd^ MIC). TXA11114 potentiated most antibiotics in a concentration-dependent manner. Tetracyclines such as doxycycline and minocycline were potentiated by up to 8- and 16-fold, respectively. All cephalosporins tested were potentiated moderately to 4-fold while aztreonam, a monobactam, was potentiated only by 2-fold. Chloramphenicol and cotrimoxazole were potentiated up to 4-fold and 8-fold respectively. As anticipated for EPIs, imipenem and gentamicin, which are not the substrate of RND efflux pumps in *P. aeruginosa*, were not potentiated by TXA11114 at any of the concentrations tested. The experimentally observed lowest active concentration for TXA11114 was 6.25 µg/mL (1/16^th^ MIC). Thus, a subinhibitory concentration of 6.25 µg/mL, unable to exert any antimicrobial effect by itself, was chosen for future antimicrobial potentiation assays in *P. aeruginosa*.

**Table 2.** Potentiation of different antibiotics classes by TXA11114 against *P. aeruginosa* ATCC 27853.

| Antibiotics | MICs (µg/mL) |  | Fold MIC reduction | Antibiotics | MICs (µg/mL) |  | Fold MIC reduction |
| --- | --- | --- | --- | --- | --- | --- | --- |
|  | No EPI | + 6.25 µg/mL TXA11114 |  |  | No EPI | +6.25 µg/mL TXA11114 |  |
| Aztreonam | 8 | 4 | 2 | Imipenem<sup>#</sup> | 4 | 4 | 1 |
| Cefepime | 2 | 0.5 | 4 | Levofloxacin | 1 | 0.125 | 8 |
| Cefpirome | 8 | 2 | 4 | Cotrimoxazole | 256 | 32 | 8 |
| Ceftazidime | 2 | 0.5 | 4 | Doxycycline | 32 | 4 | 8 |
| Ceftobiprole | 8 | 2 | 4 | Minocycline | 32 | 2 | 16 |
| Gentamicin<sup>#</sup> | 2 | 2 | 1 | Chloramphenicol | >256 | 64 | >4 |

### 4. TXA11114 outperforms known EPIs and potentiates levofloxacin in *P. aeruginosa* clinical isolates obtained from CDC-FDA and Walter-Reed Army Hospital

In a comparative study, we evaluated the levofloxacin potentiation effect of TXA11114 along with well-known EPIs PAβN and MC-04,124 against *P. aeruginosa* clinical isolates from the CDC-FDA isolate bank [13, 14]. TXA11114 was much more potent than PAβN and MC-04,124. It reduced the levofloxacin MIC in 90% of the CDC isolates by ≥ 8-fold at 6.25 μg/mL, while the potencies of other EPIs against the same isolates were negligible at 50 μg/mL (Table 3).

**Table 3.**
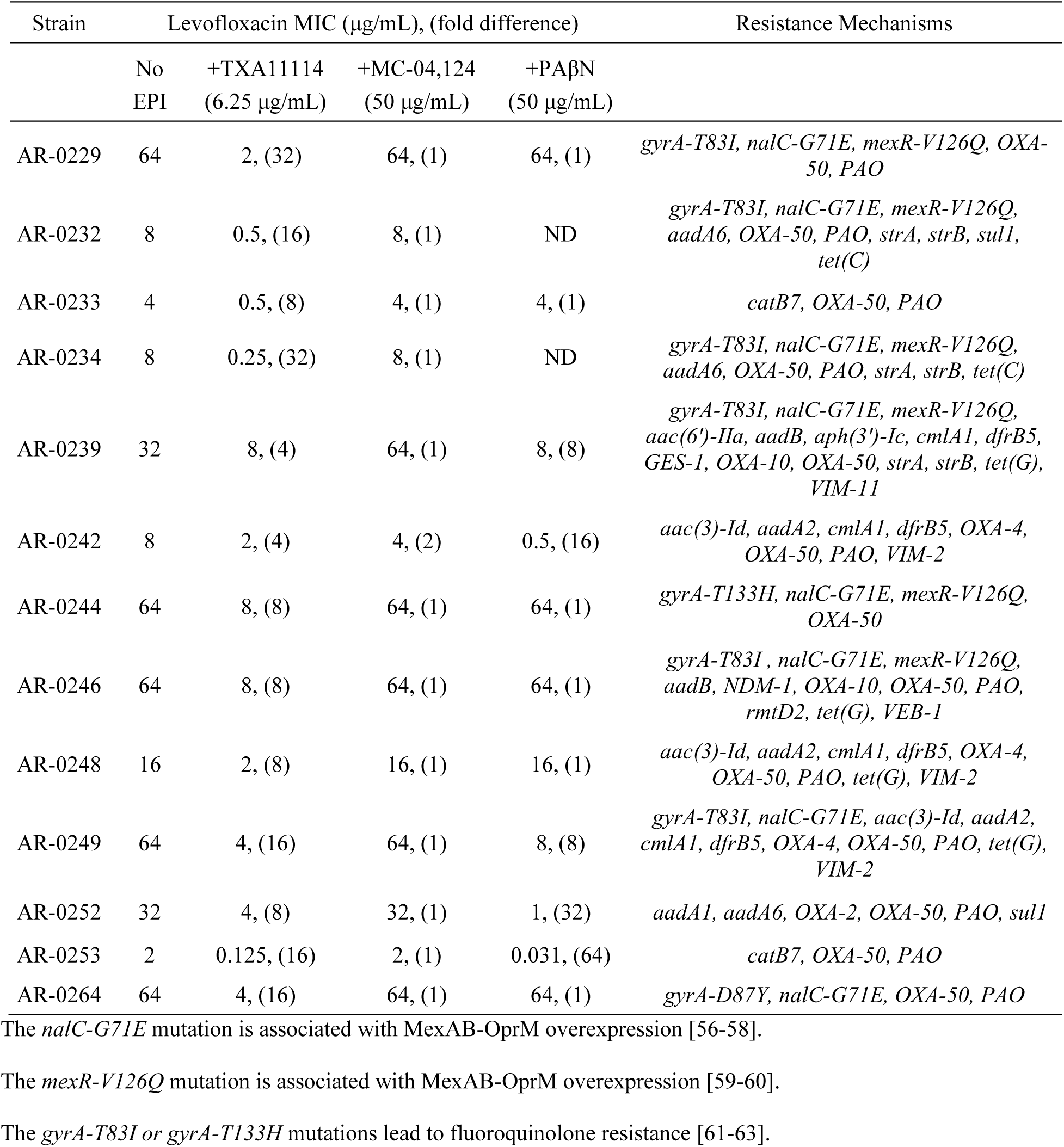
Comparative Levofloxacin potentiation study of TXA11114 and other known EPIs against multidrug-resistant *P. aeruginosa* clinical isolates from the CDC-FDA.

Similarly, when evaluated against 22 MRSN clinical isolates from the Walter Reed Army Hospital in the US [33], most of these highly MDR isolates became susceptible to levofloxacin when combined with TXA11114, while levofloxacin alone had an MIC of 8 μg/mL or above. TXA11114 at 6.25 μg/mL could potentiate all isolates 4- to 32-fold (Table 4). The resistance determinants of each of these strains are provided in Tables 3 and 4. TXA11114 could potentiate levofloxacin in more than 90% of the clinical isolates irrespective of their resistance mechanism. This further demonstrates that the TXA11114-levofloxacin combination could be highly effective in treating MDR *P. aeruginosa* infections.

**Table 4.** Levofloxacin potentiation by TXA11114 against multidrug-resistant *P. aeruginosa* Clinical Isolates from Walter Reed Army Hospital in US.

| Strains | Levofloxacin MIC (µg/mL),<br>(fold difference) | Fluoroquinolones and Efflux Resistance Mechanisms |
| --- | --- | --- |

|  | No<br>EPI | +TXA11114<br>(6.25 µg/mL) |  |
| --- | --- | --- | --- |
| MRSN #0317 | 8 | ≤0.063, (128) | *nalC-G71E, gyrA-D87N, gyrA-T133M* |
| MRSN #8130 | 16 | 2, (8) | *nalC-G71E, gyrA-D87N* |
| MRSN #994 | 16 | 2, (8) | *gyrA-T83I, mexR-V126E* |
| MRSN #1344 | 8 | 0.25, (16) | *mexR-V126E, nalC-G71E* |
| MRSN #2108 | 32 | 32, (1) | *gyrA-T83I, mexR-V126E, nalC-R209D, parS-A115E* |
| MRSN #3587 | 8 | 1, (8) | *nalC- LOF* |
| MRSN #3705 | 8 | 0.25, (32) | *nalC- LOF, parS-A115D* |
| MRSN #5498 | 32 | 4, (8) | *mexR-V126E, nalC-G71E* |
| MRSN #5519 | 64 | 4, (16) | *gyrA-T83I, mexR-V126E, parC-S87L* |
| MRSN #5539 | 16 | 8, (2) | *gyrA-T83I* |
| MRSN #6220 | 128 | 8, (16) | *gyrA-T83I, parC-S87L* |
| MRSN #6241 | 8 | 2, (4) | *mexR-V126E* |
| MRSN #6678 | 32 | 4, (8) | *gyrA-T83I, parC-S87L* |
| MRSN # 6695 | 8 | 0.5, (16) | *mexR-V126E, nalC-G71E, gyrA-T83I* |
| MRSN #8139 | 8 | 1, (8) | *nalC-G71E, gyrA-T83I* |
| MRSN #8912 | 16 | 2, (8) | *gyrA-T83I, nalC-A145V* |
| MRSN #8914 | 16 | 2, (8) | *gyrA-T83I, parE-S457G* |
| MRSN #8915 | 128 | 64, (2) | *gyrA-T83I, gyrA-D87N, parC-S87L, parS-L137P* |
| MRSN #14981 | 16 | 1, (16) | *mexR-V126E, nalC-G71E* |
| MRSN #15566 | 8 | 0.25, (32) | *mexR-V126E, nalC-G71E* |
| MRSN #16344 | 8 | 0.5, (16) | *mexR-V126E, nalC-G71E, gyrA-D87N* |
| MRSN #369569 | 32 | 4, (8) | *mexR-V126E, nalC-G71E, gyrA-T133M* |

The *nalC-G71E* mutation is associated with MexAB-OprM overexpression [56-58].
The *mexR-V126Q* mutation is associated with MexAB-OprM overexpression [59-60].
The *gyrA-T83I* or *gyrA-T133H* mutations lead to fluoroquinolone resistance [61-63].

### 5. TXA11114 does not affect outer- and inner-membrane integrity in *P. aeruginosa*

The significant toxicity of previously identified EPIs has prevented their clinical use. So, understanding the TXA11114 mechanism of action (MoA) was of paramount importance to eliminate off-target effects that may arise from other activities besides efflux pump inhibition. Historically EPIs are known to augment the action of traditional antibiotics by compromising the integrity of bacterial membranes, allowing increased drug accumulation. One of the major concerns of earlier EPIs was their ability to disrupt bacterial membranes. Two approaches were taken to investigate if membrane disruption played a role in the potentiation of antibiotics seen in Table 2 by TXA11114: (1) a flow cytometry-based propidium iodide (PI) assay to monitor inner membrane permeabilization, and (2) a nitrocefin (NCF) assay to monitor outer membrane permeabilization. NCF is a chromogenic cephalosporin that changes from yellow to red when the amide bond in the β-lactam ring is hydrolyzed by a β-lactamase. The rate of hydrolysis in intact cells is slow as it is limited by the rate of diffusion of periplasmic β-lactamase across the outer membrane. However, in the presence of an agent that permeabilizes the outer membrane, the rate of hydrolysis will increase. The impact of TXA11114 on NCF hydrolysis is minimal at concentrations below 25 μg/mL, suggesting that TXA11114 does not interact with the outer membrane of *P. aeruginosa* at these concentrations (Figure 4A). This result is comparable to what was previously reported for TXA01182 and TXA09115 [23, 24], suggesting that incorporation of the fluorinated side chain of TXA11114 did not introduce membrane disruption properties to the compound. Polymyxin B was used as a positive control (Figure 4B). In the PI assay, log-phase *P. aeruginosa* cells were mixed with various concentrations of TXA11114 followed by the addition of PI. Cells with intact membranes exclude PI and remain non-fluorescent, while cells with compromised membrane integrity allow PI to enter and bind to DNA, resulting in fluorescence. TXA11114 did not disrupt the bacterial inner membrane below concentrations of 50 μg/mL compared with water and polymyxin B as vehicle and positive controls, respectively (Figure 4C).

**Figure 4.**
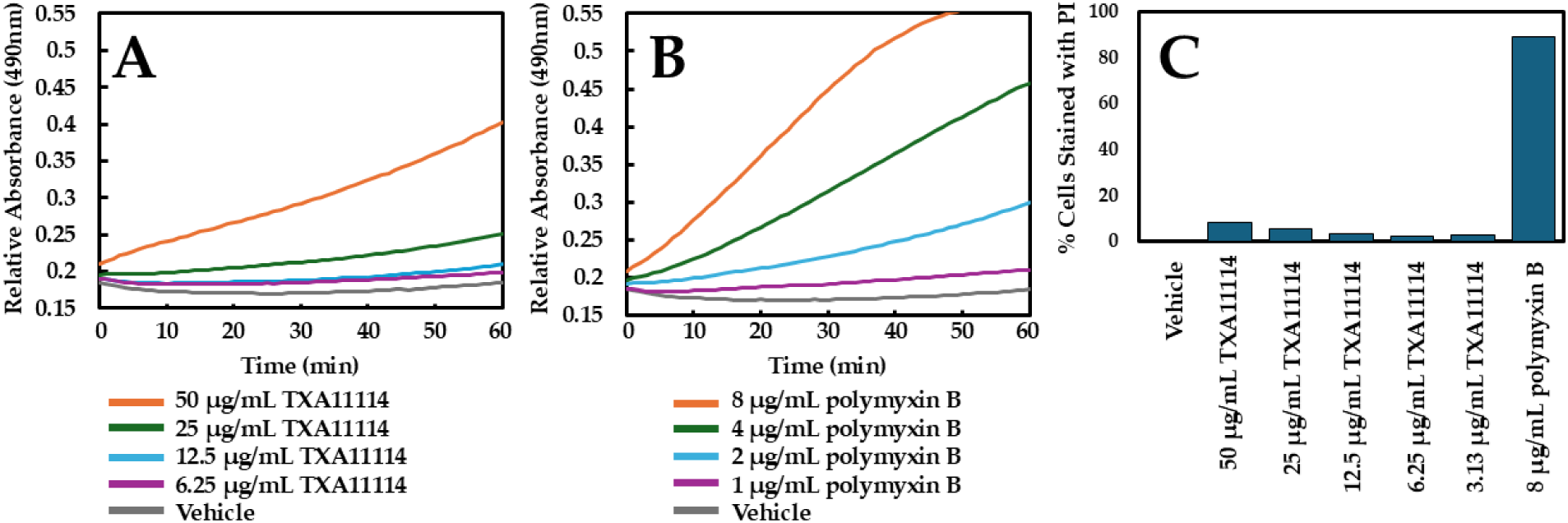
*P. aeruginosa* ATCC 27853 outer- and inner-membrane permeabilization studies with TXA11114. Hydrolysis of NCF (**A** and **B**) was used as read out for outer-membrane activity while inner-membrane activity was determined by measurement of PI fluorescence (**C**) using flow cytometry. Experiments were repeated three times to ensure reproducibility.

### 6.TXA11114 blocks efflux of ethidium bromide and levofloxacin

The ability of TXA11114 to inhibit bacterial efflux was studied using two quantitative assays. In the first assay, the efflux of ethidium bromide (EtBr) by *P. aeruginosa* cells is studied in the presence of different concentrations of TXA11114. Type strain *P. aeruginosa* ATCC 27853 cells were incubated with EtBr to allow for intracellular accumulation and treated with carbonyl cyanide 3-chlorophenylhydrazone (CCCP) to inhibit active efflux. When bound to intracellular bacterial DNA, EtBr fluoresces brightly, while any unbound EtBr outside bacterial cells exhibit little or no fluorescence. Following activation by the addition of glucose, the efflux of EtBr can be followed in real time as a decrease in fluorescence based on the concentration of TXA11114. As seen in Figure 5A, the fluorescence intensity increased proportionally with the increasing concentration of TXA11114, indicating intracellular accumulation of EtBr and supporting a role in efflux inhibition by TXA11114.

**Figure 5.**
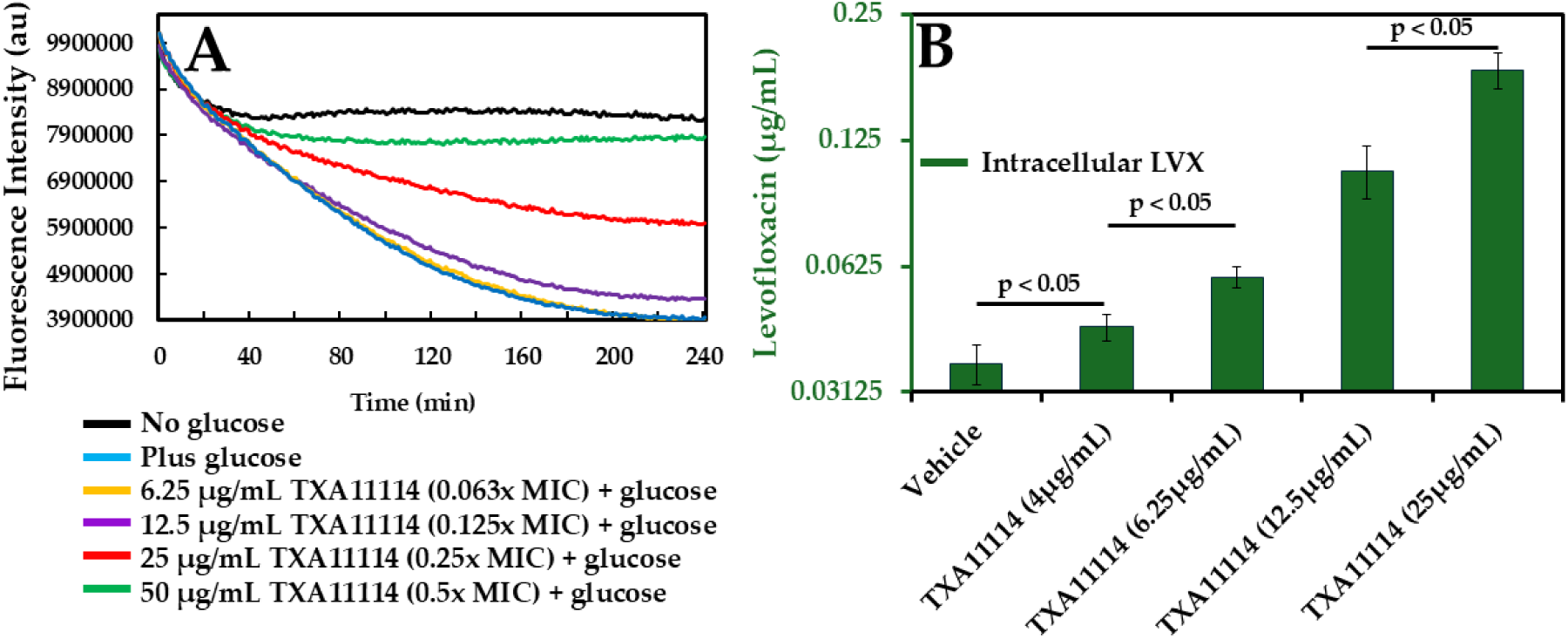
*P. aeruginosa* efflux inhibition by TXA11114. TXA11114 concentration dependent blockade of EtBr efflux (**A**) and accumulation of levofloxacin in live cells (**B**). Experiments were done in triplicates. Data is presented either as a representative value (**A**) or as a mean value with standard deviations (**B**).

The second assay measures levofloxacin accumulation inside *P. aeruginosa* following treatment with varying concentrations of TXA11114 [24], providing more direct evidence of its effectiveness as an EPI in live bacterial cells. *P. aeruginosa* DA7232 harboring mutations in DNA gyrase (*gyrA*-T83I) and topoisomerase IV (*parC*-S80L) was used for this study since it is highly resistant to levofloxacin [34]. Moreover, the levofloxacin MIC in this strain goes from 256 µg/mL to 1 µg/mL in the presence of 6.25 µg/mL TXA11114 making this strain ideal for the study. In this assay, *P. aeruginosa* DA7232 cells are incubated with levofloxacin and TXA11114 to allow for intracellular accumulation. After washing and membrane permeabilization, intracellular levofloxacin accumulation was monitored by following the changes in levofloxacin fluorescence. Similar assay was described elsewhere for the same purpose [35–36]. TXA11114 led to the accumulation of levofloxacin inside *P. aeruginosa* in a concentration-dependent manner is in support of its role in efflux inhibition.

### 7. TXA11114 shows no effect on the inner membrane potential and cellular ATP content

RND efflux pumps require an active proton gradient across the inner membrane to flush out antimicrobial molecules [37]. Thus, disruption of the membrane potential in bacteria may also disable RND efflux function and lead to antimicrobial potentiation. Whether TXA11114 treatment dissipated the membrane potential in *P. aeruginosa* was investigated with the fluorescence molecular probe 3,3’-diethyloxacarbocyanine iodide (DiOC_2_(3)) [38]. DiOC_2_(3) accumulates within cells with polarized membranes, shifting from green to red fluorescence at high concentration due to concentration-dependent dye stacking [39]. Membrane depolarization prevents this shift. As shown in Figure 6A, treatment of *P. aeruginosa* with TXA11114 at concentrations ranging from 20 to 0.01 μM (20 μM = 7.32 µg/mL TXA11114) did not lead to significant membrane depolarization ruling out membrane depolarization as the MoA responsible for antibiotic potentiation. Azithromycin and CCCP were used as positive and negative controls, respectively.

**Figure 6.**
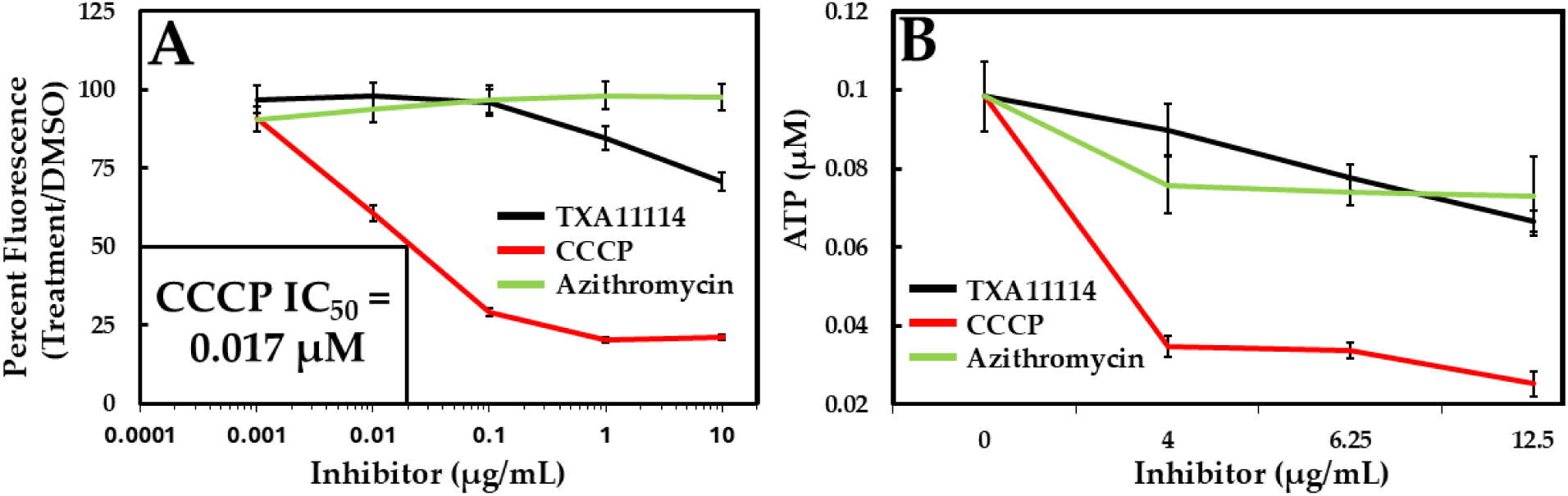
Bacterial membrane potential was assessed by measuring fluorescence changes in the membrane potential indicator DiOC_2_(3) (**A**). Cellular ATP pool was measured by luminescence-based assay (**B**). Experiments are done in triplicates; data is presented as mean value with standard deviations (**B**).

Further, a deviation from normal membrane function can affect the activity of respiratory components and diminish ATP synthesis. Thus, an indirect approach to assess a drug’s membrane effect is by monitoring cellular ATP levels after treatment. To exclude ATP depletion as a TXA11114 MoA, bacterial ATP levels were evaluated three hours post treatment. As shown in Figure 6B, *P. aeruginosa* treatment with TXA11114 did not result in ATP depletion. In contrast, significant depletion of ATP was observed in CCCP treated cells, compared to untreated control. Azithromycin was used as a negative control.

### 8. TXA11114 potentiates levofloxacin in efflux pump overexpressed strains

Next, the ability of TXA11114 to inhibit *P. aeruginosa* mutants overexpressing MexAB-OprM was evaluated. Loss of function mutations in *nalB* resulting in overexpression of MexAB-OprM, have been identified in *P. aeruginosa* clinical isolates and are associated with resistance to cephalosporins, fluoroquinolones and aminoglycosides [40–46]. In theory, an EPI should reverse the antimicrobial susceptibility lost in bacteria overexpressing efflux pumps. Thus, the TXA11114 EPI activity was studied using antibiotics that display efflux liabilities in this RND efflux pump (Table 5). The level of antibiotic potentiation seen with TXA11114 in MexAB-OprM-overproducing strain (K1455) was 2- to 4-fold higher than the parent strain (K767). In agreement with being an EPI, TXA11114 did not significantly potentiate any antibiotic in the in MexAB-OprM deficient strain (K1119). Finally, imipenem, which is not the substrate of RND efflux pumps in *P. aeruginosa*, was not potentiated by TXA11114 in any of the strains tested.

**Table 5.** Comparative study of antibiotic MIC potentiation by TXA11114 in efflux mutants.

| Antibiotics | MIC ratio of Antibiotics alone/(Antibiotics +<br>6.25 µg/mL TXA11114) |  |  |
| --- | --- | --- | --- |
| | K767<br>(WT) | K1455<br>($\uparrow mexAB-oprM$) | K1119<br>($\Delta mexAB-oprM$) |
| Levofloxacin | 8 | 16 | 2 |
| Doxycycline | 8 | 32 | 2 |
| Chloramphenicol | 4 | 16 | 2 |
| Cefepime | 8 | 16 | 2 |
| Ceftazidime | 4 | 8 | 2 |
| Imipenem<sup>#</sup> | 1 | 1 | 1 |

<sup>#</sup> Not the substrates for efflux pumps in *P. aeruginosa*.

### 9. TXA11114 prolonged the levofloxacin post antibiotic effect

Antimicrobials exhibit a post antibiotic effect (PAE) after their removal and introduce a growth delay on bacterial cultures when compared to the untreated condition. An exponentially grown *P. aeruginosa* ATCC 27853 was treated with 1x MIC of levofloxacin in the presence and absence of 6.25 µg/mL of TXA11114 for 1 hour. After treatment, drug concentrations were reduced by diluting cultures 50-fold, and bacterial growth was monitored by measuring optical density. When compared to untreated control, bacterial cultures treated with levofloxacin alone displayed a PAE of 4.9 hours, whereas TXA11114-levofloxacin combination treated bacteria displayed a PAE of 6.3 hours. Thus, the combination prolonged the levofloxacin PAE by 1.4 hours. The prolonged PAE of the combination is consistent with EPI-driven levofloxacin accumulation within the cell and could aid in determining the optimum dosing frequency of the antibiotic by ensuring properly spaced dosing intervals.

### 10. TXA11114-levofloxacin combination has undetectable level of resistance in drug-sensitive and drug-resistant strains

In addition to reducing the levels of intrinsic resistance, a potent EPI is also expected to significantly reverse acquired resistance as well as decrease the frequency at which antimicrobial resistance evolves. A resistance study for the TXA11114-levofloxacin combination was investigated in drug-sensitive (ATCC 27853) and drug-resistant (AR-0232, AR-0248 and AR-0249) *P. aeruginosa* strains. We found that the combination resulted in undetectable levels of spontaneous levofloxacin resistance against 10^10^ CFU/mL bacterial inputs. On the contrary, standalone levofloxacin resulted in significant levels of resistance from the same bacterial input (10^10^ CFU/mL, Table 6). The undetectable levels of resistance highlight the clinical utility of these combinations to limit the selection of spontaneous resistant mutants, particularly in cystic fibrosis patients infected with *P. aeruginosa*, where patients are colonized by hypermutable strains that persist for years [47].

**Table 6.**
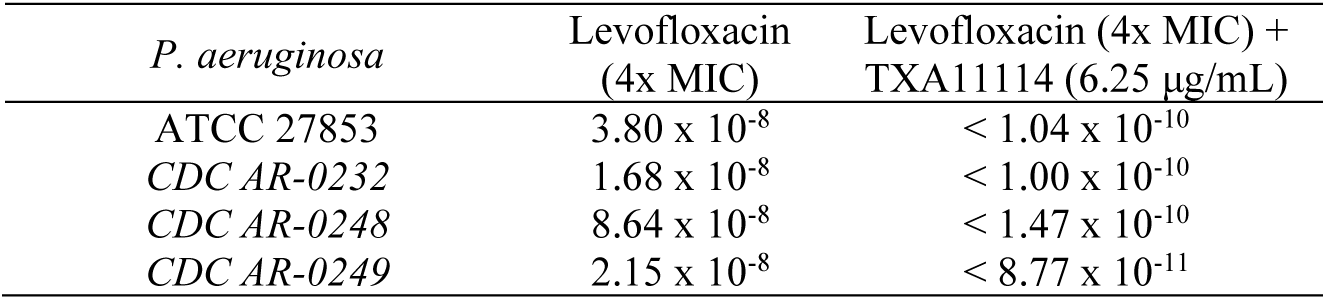
Frequency of resistance to TXA11114 and levofloxacin.

**Table 7.**
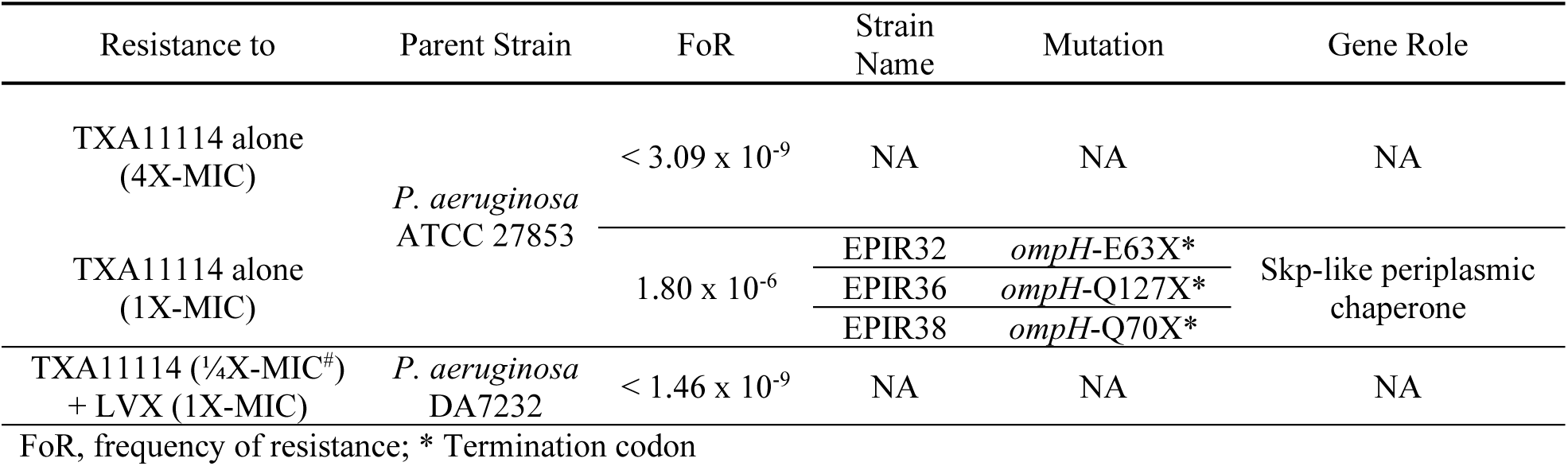
**Mutations involved in TXA11114 resistance in *P. aeruginosa***

### 11. Genetic study of TXA11114 resistance in *P. aeruginosa*

A study was conducted with the goal of understanding how *P. aeruginosa* becomes resistant to TXA11114, alone or in combination with levofloxacin. To understand resistance to TXA11114 alone, *P. aeruginosa* ATCC 27853 was exposed to 4-times or 1-time the TXA11114 MIC (400 or 100 μg/mL, respectively). All resistant colonies were verified by determining the TXA11114 MIC. Resistance to TXA11114 did not result in cross-resistance to other known antibiotics (Supplemental Table 1). No resistance was detected to 4-times the MIC of TXA11114 alone. Resistance to 1-time the MIC of TXA11114 arose at a frequency of 1.80 x10^-6^ and resulted in single point mutations that introduced early stop codons in the *ompH* gene (strains EPIR32, EPIR36 and EPIR38). OmpH is a homolog of the outer membrane chaperone Skp [48] and is overexpressed in *P. aeruginosa* isolates resistant to ampicillin and kanamycin [49]. Deletion of *ompH* results in hyper susceptibility to ertapenem, ceftazidime, levofloxacin, tigecycline, and cotrimoxazole [50]. Consistent with this, EPIR32-38 exhibited increased susceptibility to levofloxacin, ceftazidime, tigecycline, doxycycline, meropenem, and amikacin (Supplemental Table 1). Notably, the increased susceptibility to levofloxacin and doxycycline in some of these mutants align with the potentiation observed at the lowest active concentration of TXA11114 in the parent strain (Table 2). In *E. coli*, Skp interacts with over 30 envelope proteins [51, 52], and *ompH* genetically interacts with *acrD*, a component of the AcrAD-TolC aminoglycoside efflux pump [53, 54]. Since AcrD shares 62% sequence similarity with *P. aeruginosa* MexB, OmpH may assist in folding an efflux pump component targeted by TXA11114. If so, OmpH loss could lead to misfolded efflux pumps, reducing antibiotic efflux and increasing susceptibility. Alternatively, TXA11114 may directly inhibit OmpH, disrupting efflux pump folding and enhancing strongly antibiotic susceptibility.

### 12. TXA11114 promotes bacterial killing by levofloxacin

In addition to assessing the potentiation activity of TXA11114 *in vitro,* its potentiation of a minimally bactericidal concentration of levofloxacin (1X-MIC) was probed against *P. aeruginosa* ATCC 27853 and AR-0232 with time-kill studies. Figure 7 shows time-kill curves with levofloxacin alone or combined with TXA11114. By itself, TXA11114 had no effect on the growth of *P. aeruginosa* strains at 6.25 µg/mL (dark green curve). TXA11114 enhanced levofloxacin killing kinetics in a concentration-dependent manner (light green, purple and light blue curves). The combinations reduced bacterial counts by more than 3.42 log₁₀ after 3 hours, 1.68 log₁₀ after 6 hours, and 7.97 log₁₀ after 24 hours compared to levofloxacin alone. These results suggest that the killing kinetics for the TXA11114/levofloxacin combination are faster than those of levofloxacin alone, similar to what was reported previously for the TXA01182/levofloxacin and TXA09155/moxifloxacin combinations [23, 24].

**Figure 7.**
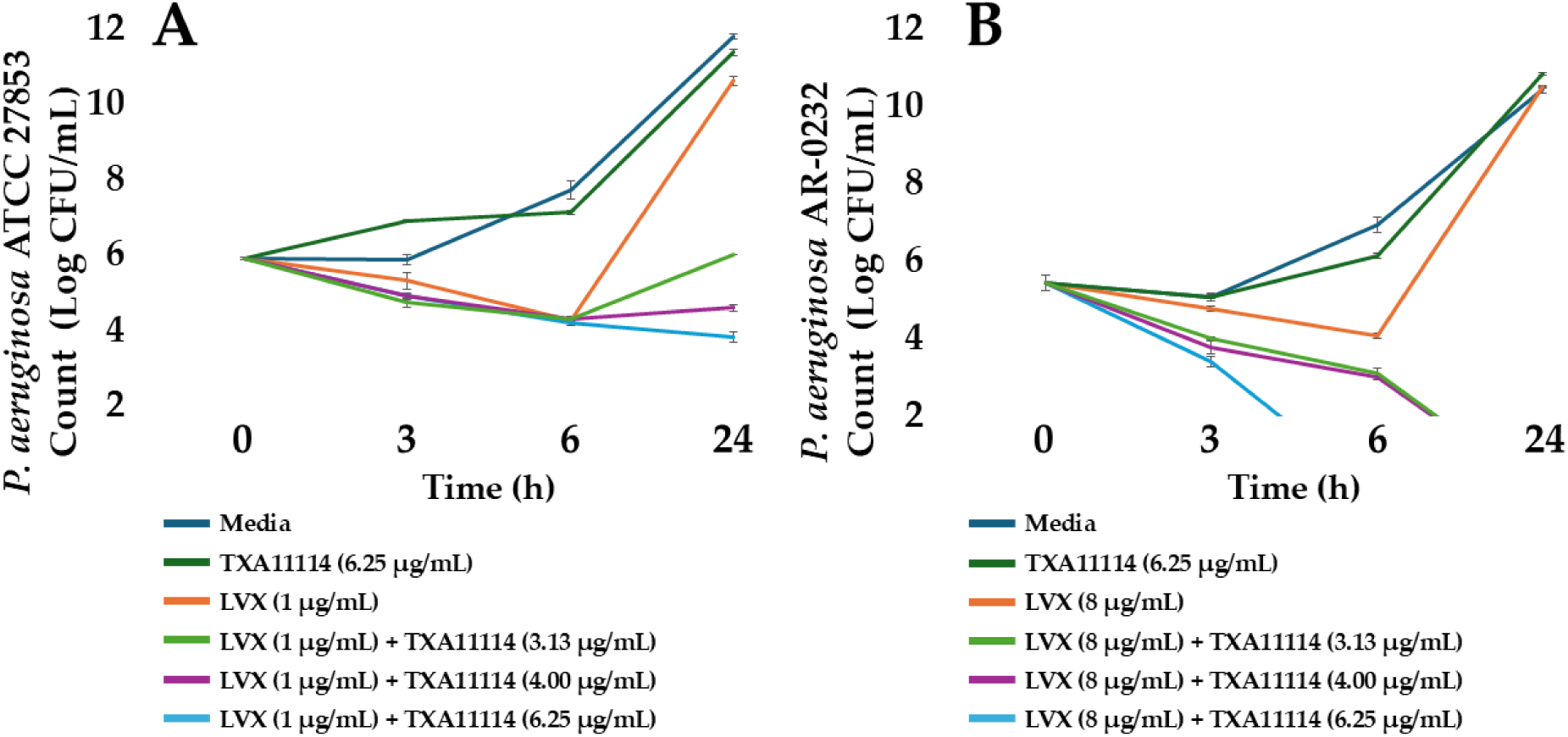
Time killing kinetics of levofloxacin alone or in combination with different doses of TXA11114 against *P. aeruginosa* type strain ATCC 27853 (**A**) and clinical isolate AR-0232 (**B)**. Data is presented as mean values with standard deviations derived from technical triplicates.

### 13. TXA11114 has good physiochemical and ADME properties

Along with its microbiological evaluation, TXA1114 was evaluated for its physiochemical and *in vitro* ADME properties. In general, TXA11114 follows Lipinski’s rule-of five, a rule describing molecular properties believed to be important for a drug’s pharmacokinetics in the human body, in addition to being highly soluble (>145 µM at pH 7.4). Table 8 summarizes the *in vitro* chemical absorption, distribution, metabolism, excretion, and toxicity (ADMET) profile of TXA11114. Overall, the plasma stability of TXA11114 is good across four species (human, dog, rat and mouse) with t_1/2_ exceeding 2h. The microsomal stability across two species (human and rat) are equally good, with t_1/2_ exceeding 1h. It did not inhibit cytochrome P450 isoforms having IC_50_ values greater than 100 µM for CYP1A2, CYP2C19, CYP2C9, and CYP2D6 while it is 63 µM for CYP3A4.

**Table 8.** TXA11114 has favorable Physicochemical and ADME properties.

|  |  |  |  | Human | Dog | Rat | Mouse |  |
| --- | --- | --- | --- | --- | --- | --- | --- | --- |
| Protein Binding (%) |  |  |  | 99 | 98.9 | 98.5 | 98.9 |  |
| Plasma stability t<sub>1/2</sub> (min) |  |  |  | >120 | >120 | >120 | >120 |  |
| Microsome Stability t<sub>1/2</sub> (min) |  |  |  | >60 | ND | >60 | ND |  |
| MW | Aqueous solubility (μM) at pH7.4 | cLogP | PSA | CYP Inhibition IC50 (μM) |  |  |  |  |
|  |  |  |  | 1A2 | 2C9 | 2C19 | 2D6 | 3A4 |
| 372.42 | 145.9 ± 6.1 | 2.34 | 93.17 | >100 | >100 | >100 | >100 | 63 |

The plasma protein binding in the four species is between 98-99%. The observed cytotoxicity IC_50_ values of TXA11114 in CellTiterGlo^TM^ assay (293T, A549) were all greater than 20 µM.

### 14. TXA11114 has a low cardiotoxicity potential

Unwanted binding to voltage-gated ion channels in eukaryotic cells can lead to significant side effects due to their role in pharmacokinetics and pharmacodynamics [55]. To see how the strategic placement of fluorine in the TXA11114 diamine side chain affected the activities on the three ion channel targets (voltage-gated sodium: Nav1.5 (peak), voltage-gated potassium: hERG, and voltage-gated calcium: Cav1.2) was assayed using Eurofins’ QPatch electrophysiological platform (Table 9). IC_50_ values were determined by a non-linear, least squares regression analysis. Reference standards were run as an integral part of each assay to ensure the validity of the results obtained. Results showing an inhibition higher than 50% are considered to represent significant effects of test compound. As shown in Table 9, TXA11114 (10 µM) has −3.7% inhibition of hNav1.5, 0.2% inhibition of hERG and 10.9% inhibition of hCa1.2. These results suggest that TXA11114 might have low potential for cardiotoxicity and other side effects like neurological disturbances, muscle weakness, altered sensory perception, and seizures or respiratory depression.

**Table 9.**
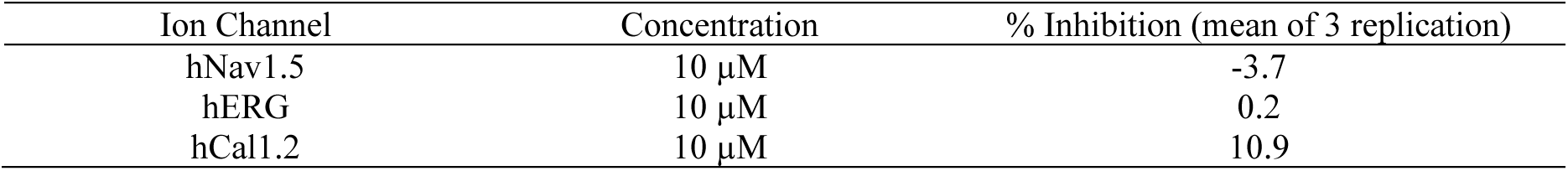
CiPA Core Panel Data for TXA11114 (cardiotoxicity monitoring)

### 15. TXA11114 displays minimal nephrotoxicity

As mentioned before, nephrotoxicity due to the intralysosomal accumulation of basic molecules in renal tissue is a concern since our EPIs contain two primary amine groups [27]. To investigate if the introduction of fluorine in the diamine side chain helped in reducing any possible nephrotoxicity, we evaluated TXA11114 induced nephrotoxicity by measuring blood urea nitrogen (BUN) and concentration of serum creatinine (CRE). BUN and CRE tests were conducted by SRI International (Menlo Park, CA) to evaluate potential nephrotoxicity following TXA11114 administration. Briefly, adult male Sprague Dawley rats (3 rats/group) were treated with TXA11114 (10 mg/kg, IV, QD) and blood samples were collected 24 hours after dosing. Table 10 shows BUN and CRE levels following TXA11114 treatment. BUN and CRE levels following administration fell within normal historical ranges and were considered of minimal toxicologic significance, the elevated level of which may implicate renal toxicity. These results suggest that TXA11114 administration did not cause nephrotoxicity in rats.

**Table 10.**
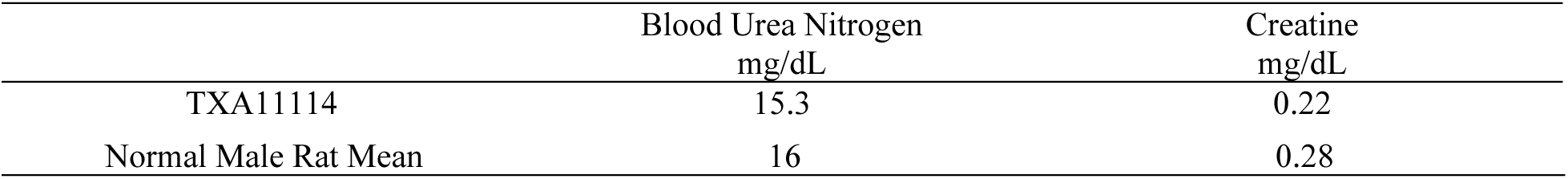
Nephrotoxicity assessment for TXA11114.

### 16. TXA11114 and levofloxacin have complementary pharmacokinetic profile

To provide the maximum pharmacodynamic (PD) benefit, the pharmacokinetics (PK) of an EPI should be complimentary to the PK of the antibiotic component of the combination. To see if our EPI class has complimentary PK properties with the partner antibiotic levofloxacin, plasma and bronchoalveolar lavage fluid (BALF) PK levels were determined for TXA11114 w/wo levofloxacin in *P. aeruginosa* lung infected mice (Figure 8A and 8C). Likewise, PK levels were determined for levofloxacin w/wo TXA11114 (Figure 8B and 8D). Overall, plasma levels of TXA11114 were comparable when dosed by itself or co-administered with levofloxacin. At 30 mg/kg, TXA11114 exhibited a peak concentration of 12.98 – 16.54 µg/mL (at 0.083 hr), total plasma exposure of (AUC) of 30.7 – 33.1 µg-hr/mL, 2.42 – 2.50 L/kg volume of distribution, 0.87 – 0.95 L/hr/kg plasma clearance and elimination half-life of 1.84 – 1.93 hr (Figure 8E). Plasma levels for levofloxacin were also comparable when administered alone or in combination with TXA11114. At 30 mg/kg, levofloxacin exhibited a peak concentration of 24.1 – 24.39 µg/mL (at 0.25 hr), total plasma exposure of (AUC) of 32.9 – 42.3 µg-hr/mL, 1.59 – 1.89 L/kg volume of distribution, 0.68 – 0.89 L/hr/kg plasma clearance and elimination half-life of 1.48 – 1.63 hr. PK levels in BALF samples correspond to analysis of the fluid itself and have not been corrected for the dilution of epithelial lining fluid (based on a urea assay, Figure 8E). Slight variations in TXA11114 levels were observed at each time point sampling for both dose groups. Overall lung exposures were similar with C_max_ concentrations of 0.75 – 1.06 µg/mL observed at 4 – hrs and AUC values of 12.3 – 13.4 µg-hr/mL. Measured levofloxacin peak levels and overall exposure (AUC) were 2.0 & 2.56 µg/mL and 3.9 & 6.3 µg-hr/mL for animals administered levofloxacin alone and levofloxacin with TXA11114, respectively (Figure 8E). These results suggest that administration of TXA11114 or levofloxacin alone are comparable, and that administration of the combination did not disrupt the exposure levels of the individual components.

**Figure 8.**
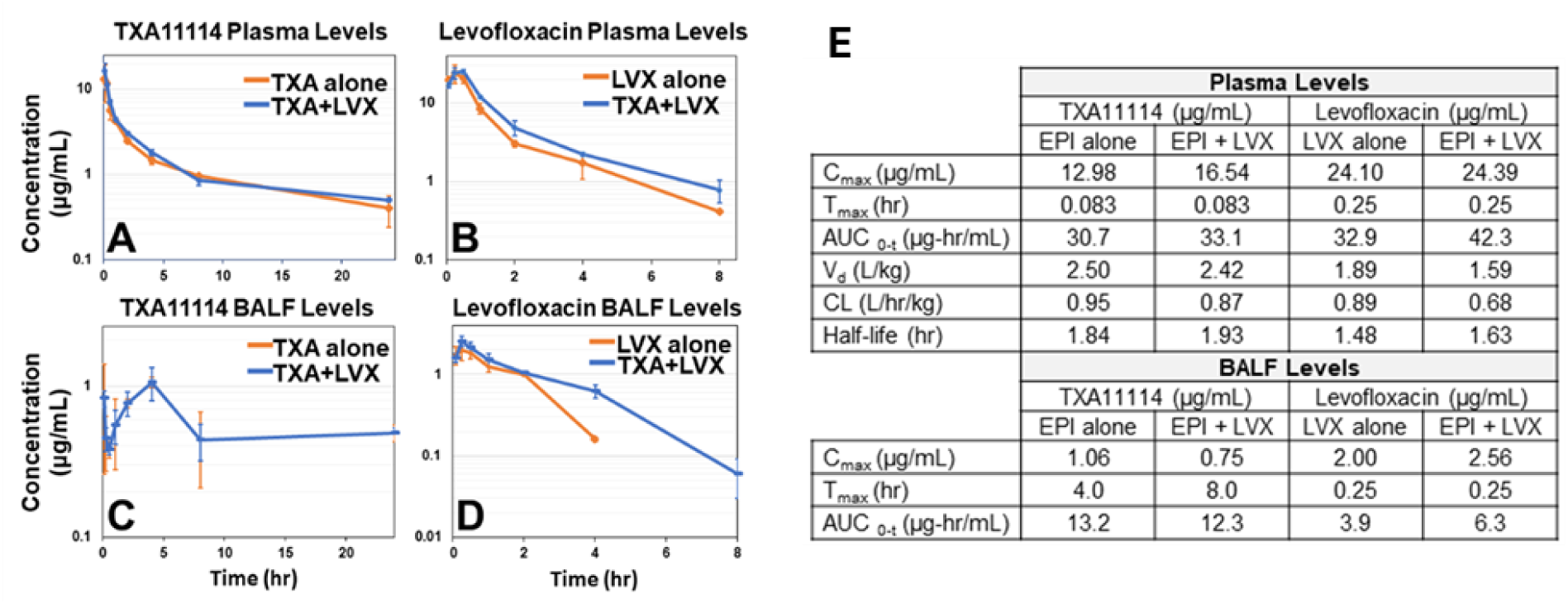
Pharmacokinetic studies of TXA11114

### 17. TXA11114 has improved acute toxicity profile

With overall acceptable microbiological, physiochemical, ADME, toxicity and complementary pharmacokinetic characteristics, the next task was to evaluate TXA11114’s *in vivo* efficacy in a *P. aeruginosa* infection model. Thus, the safety and tolerability of TXA11114 was determined to find an optimum dosing capability. The maximum tolerated dose (MTD) of TXA11114 was assessed in female mice (CD-1, 6-8 weeks old) at University of North Texas Health Science Center (UNTNHS). TXA11114 was administered IV to groups of three female mice. Mice are administered TXA11114 in the range 30-240 mg/kg. The first (30 mg/kg) dose level was administered and mice observed for any effects (including respiration, piloerection, startle response, skin color, injection site reactions, hunched posture, ataxia, salivation, lacrimation, diarrhea, convulsion, death and others if observed) for approximately 10 minutes before proceeding to the next higher dose. As doses are tolerated (not resulting in effects such as mortality, convulsions, or other severe morbidity), they are increased. Survival and general observations as to the tolerability of the administered dose started immediately and continued for a period of 48 hours after each dose was recorded. Following this dosing regimen, an MTD for TXA11114 was determined to be greater than 60 mg/kg and less than 90 mg/kg without causing any acute toxicity. For comparison, the non-fluorinated EPI TXA01182 has an MTD of 12.5 mg/kg while the constrained EPI TXA09155 has an MTD of 30 mg/kg. Clearly the substitution of fluorine in the diamine side chain improved the MTD by more than 2- to ∼5-fold as compared to the non-fluorinated analogs.

### 18. TXA11114 and levofloxacin combination is efficacious in murine thigh and lung infection models

The translation of *in vitro* potency to *in vivo* efficacy in animal infection models remains a major challenge in the development of EPIs. We evaluated TXA11114’s ability to enhance levofloxacin’s antibacterial activity against *P. aeruginosa* ATCC 27853 in murine thigh and lung infection models, following the protocols detailed in the supporting information.

In the thigh infection model, mice were inoculated intramuscularly (IM) in the right thigh with 5.75 log₁₀ CFU. Mean bacterial thigh titers reached 6.08 log₁₀ CFU at 2 hours and 8.82 log₁₀ CFU at 24 hours post-infection, confirming successful colonization. TXA11114 (30 mg/kg, IV, QID) alone had no effect on bacterial burden (Figure 9A). Levofloxacin alone (10, 15, and 20 mg/kg, SC, QID) exhibited concentration-dependent activity. However, the combination of TXA11114 (30 mg/kg, IV, QID) with levofloxacin (10, 15, or 20 mg/kg, SC, QID) significantly reduced bacterial burden beyond levofloxacin alone (p = 0.033 and 0.059 for the two highest combinations, Figure 9A).

**Figure 9.**
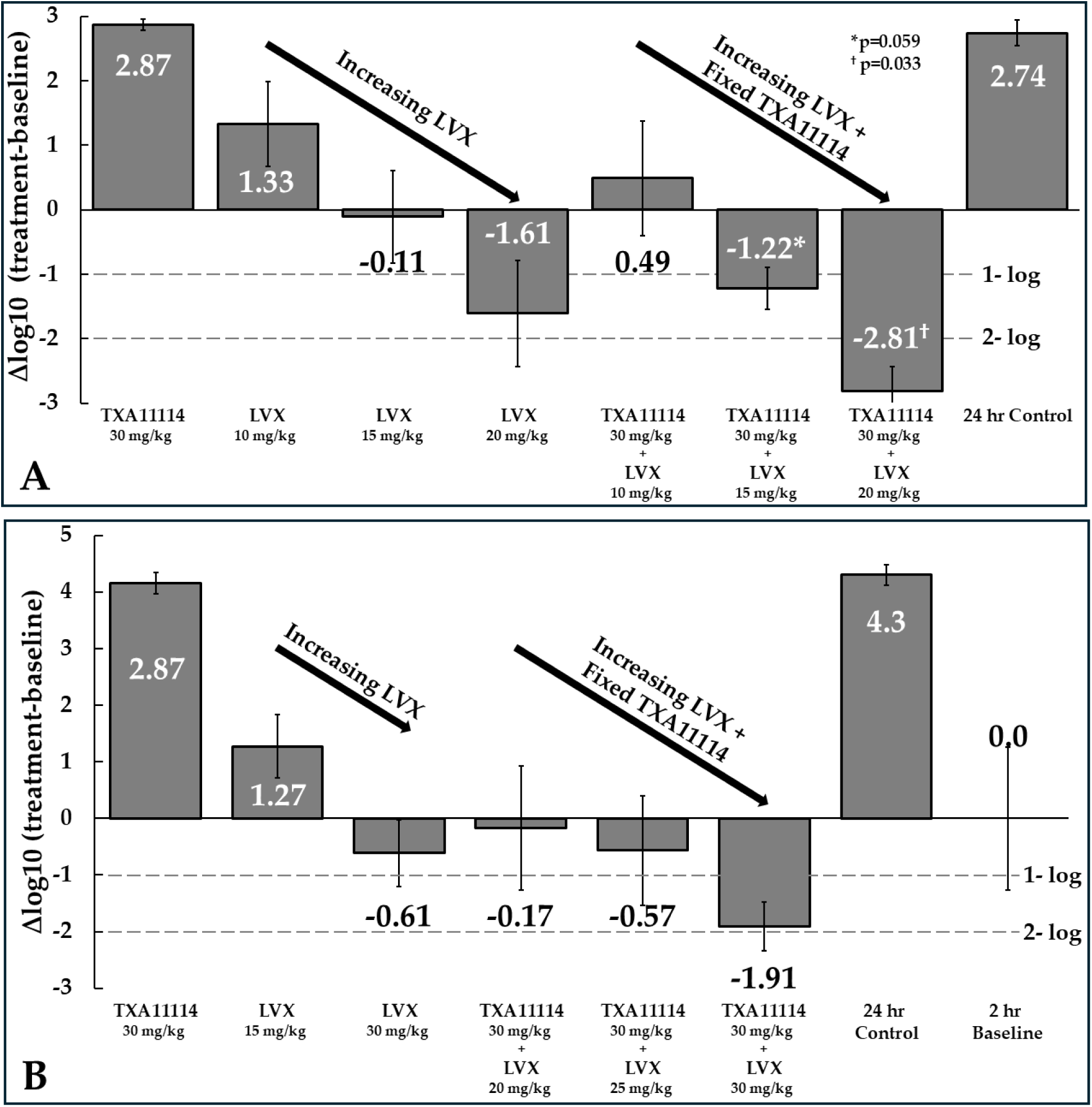
*In vivo* efficacy in murine thigh (**A**: upper) and lung (**B**: lower) models.

In the lung infection model, mice were inoculated intranasally with 4.83 log₁₀ CFU of *P. aeruginosa* ATCC 27853. Mean lung bacterial titers were 4.59 log₁₀ CFU at 2 hours and 8.89 log₁₀ CFU at 24 hours post-infection. TXA11114 (30 mg/kg, IV, QID) alone had no effect on lung bacterial counts. Levofloxacin alone (15 and 30 mg/kg, SC, QID) showed concentration-dependent activity. The combination of TXA11114 with levofloxacin resulted in a dose-dependent reduction in bacterial burden (Figure 9B). Notably, the 30 + 30 mg/kg TXA11114 + levofloxacin group reduced bacterial counts by 1.3 log₁₀ CFU more than the corresponding levofloxacin-alone group (Figure 9B).

These findings demonstrate that the complementary PK properties of TXA11114 and levofloxacin translate into enhanced *in vivo* efficacy, with the combination reducing bacterial burden by more than 1-log₁₀ below the 2-hour baseline in both thigh and lung infection models.

## Conclusion

In pursuit of a safe and effective EPI to establish this adjuvant strategy as a viable therapeutic option for treating *P. aeruginosa* infections, we designed, synthesized, and evaluated fluorine-substituted diamine-linked 2-carboxy amide indole analogs. Of the three fluorinated analogs, only TXA11114 exhibited activity, while the difluoro analog was completely inactive, likely due to its significantly reduced amine pKa, which may have impacted its activity. Although TXA11167 had a pKa comparable to TXA11114, it was unexpectedly inactive as a levofloxacin adjuvant, suggesting that factors beyond pKa influence activity.

Further microbiological screening of TXA11114 revealed a favorable profile, including strong potentiation against multiple MDR strains, rapid bactericidal activity, and an undetectable frequency of resistance when combined with levofloxacin. Biophysical and genetic studies confirmed that its mechanism of action is through efflux pump inhibition, ruling out membrane disruption or other modes of action.

Additionally, TXA11114 demonstrated an acceptable physicochemical and toxicity profile, enhancing its potential as a safe EPI. Most importantly, it exhibited a complementary pharmacokinetic profile with levofloxacin, leading to robust *in vivo* efficacy in both murine thigh and lung infection models. Given its strong potential, ongoing studies are evaluating pure diastereomeric analogs with variations in amine and fluorine stereocenters. The findings from these investigations will be reported in due course.

## Acknowledgements

This research is supported by the Cooperative Agreement Number IDSEP160030 from ASPR/BARDA, by awards from Welcome Trust and Germany’s Federal Ministry of Education and Research. The research reported in this publication was also supported by the National Institute of Allergy And Infectious Diseases of the National Institutes of Health under Award Number R44AI174351. The contents are solely the responsibility of the authors and do not necessarily represent the official views of the HHS Office of the Assistant Secretary for Preparedness and Response or other CARB-X funders and the official views of the National Institutes of Health.

## Methods

### 1. Bacterial strains, media, and reagents

*P. aeruginosa* ATCC 27853 was obtained from the American Type Culture Collection (ATCC). *P. aeruginosa* multidrug-resistant isolates were obtained from the CDC and FDA Antibiotic Resistance Isolate Bank. *P. aeruginosa* DA7232 (*gyrA*-T83I, *parC*-S80L) was a kind gift from Prof. Dan I Andersson, Uppsala University, Uppsala, Sweden and has been characterized elsewhere [34]. *P. aeruginosa* strains K767 (WT), K1455 *(mexAB-oprM* overexpressed), K2415 (*mexXY-oprM* overexpressed) and K3698 (*oprM*Δ) were obtained from Prof. Keith Poole, Queen’s University, Kingston, Ontario, Canada and have been characterized elsewhere [7–8]. Bacterial cells were grown in cation-adjusted Mueller Hinton (CAMH) media, brain heart infusion broth (BHI) or tryptic soy agar (TSA) plates all obtained from Becton, Dickinson, and Company (BD, Franklin Lakes, NJ). Aztreonam, ceftazidime, moxifloxacin, levofloxacin, minocycline, tigecycline, chloramphenicol, nitrocefin and imipenem were purchased from TOKU-E (Bellingham, WA). Azithromycin was purchased from Tokyo Chemical Industry (Portland, OR). Cotrimoxazole was purchased from Toronto Research Chemicals (Ontario, Canada). Doxycycline, polymyxin B were purchased from Sigma-Aldrich (St. Louis, MO). Ethidium bromide (EtBr) and glucose were purchased from VWR (Radnor, PA). MC-04,124 and TXA01182 were synthesized at TAXIS Pharmaceuticals. Carbonyl cyanide 3-chlorophenylhydrazone (CCCP) was purchased from Enzo Life Sciences (Farmingdale, NY). DiOC_2_(3) and propidium iodide were obtained from Thermo Fisher Scientific (Waltham, MA, US).

### 2. Determination of minimum inhibitory concentration (MIC)

MIC assay for potentiation of antimicrobial activity against P. aeruginosa MIC assays were performed as described previously [23–24].

### 3. Flow cytometry assay for permeabilization of inner cell membranes to propidium iodide (PI)

A flowcytometry assay used for assessing potential inner membranes permeabilization of *P. aeruginosa* bacterial cells to PI was conducted using the LIVE/DEAD BacLight Kit from Invitrogen (Waltham, MA). Briefly, log-phase *P. aeruginosa* ATCC 27853 bacterial cells grown in BHI broth were diluted 5-fold in PBSM (1X PBS, 1MgCl_2_) to an approximate concentration of 6.5 × 10^7^ CFU/mL. The bacteria were aliquoted into tubes and mixed with TXA11114 at concentrations ranging from 1 to 1/8^th^ times the MIC (50 to 6.25 μg/mL). Water alone was used as a solvent control. Polymyxin B was used as a positive control. Intracellular PI fluorescence was detected by flow cytometry using a CytoFlex (Beckman Coulter Inc., Brea, CA, US). The 488 nm laser was used for excitation, with the PC5.5 and FITC channels being used for emission. For each sample, the fluorescence of 10,000 individual bacterial cells was measured, and the percent of cells that stained positive for PI fluorescence was calculated.

### 4. Nitrocefin cellular assay for outer cell membrane permeabilization

Assessment of outer membrane permeabilization in *P. aeruginosa* was carried out as described previously [23–24].

### 5. Assessment of ethidium bromide (EtBr) efflux inhibition

Efflux of EtBr from *P. aeruginosa* in the presence of TXA11114 was carried out as described previously [23–24].

### 6. Levofloxacin accumulation assay

*P. aeruginosa* DA7232 was grown to an OD_600_ of 0.6 and treated with levofloxacin (64 μg/mL=1/4^th^ MIC) alone or in combination with TXA11114 at sub-inhibitory concentrations (1/4^th^, 1/8^th^ and 1/16^th^ MIC). The bacterial culture samples were treated on ice for 15 min, centrifuged, washed with PBSMG (1X PBS, 1mM MgCl2, 100mM glucose) and then resuspended in 1 ml of glycine-HCl buffer (pH 3.0) overnight for cell lysis, followed by centrifugation. The fluorescence of 100 μL of supernatant was read at 490nm following a 355nm excitation in a SpectraMax iD5 spectrophotometer (Molecular Devices).

### 7. Membrane polarization assay

*P. aeruginosa* membrane polarization assays were conducted as described previously [23–24] using strain ATCC 27853. Briefly, mid-exponential phase (OD_600_ of 0.5) *P. aeruginosa* was pelleted by centrifugation at 3000× g for 10 min at room temperature. Cells were then resuspended in 1× PBS, treated with 10 mM EDTA for 5 min and then centrifuged at 300 × g for 10 min to remove EDTA. EDTA-treated cells were pelleted and resuspended to an OD_600_ of 1.0 in assay resuspension buffer [23–24]. A 6 mM DiOC_2_(3) stock in DMSO was added to cells for a final concentration of 30 μM. DiOC_2_(3)-loaded cells were then added to a 96-well black bottom microplate for a final volume of 200 μL. TXA11114 or control compounds were added to the bottom of the well of the microplate prior to the addition of the DiOC_2_(3)-loaded cells. After 15 min incubation at 37^0^C in dark, DiOC_2_(3) fluorescence was recorded using the SpectraMax iD5 spectrophotometer (Molecular Devices) using 450-nm excitation. Red fluorescence intensity was recorded at 670-nm emission.

### 8. Determination of intracellular ATP levels

*P. aeruginosa* intracellular ATP levels were determined as described previously [23–24] using an ATP determination kit (Invitrogen, Life Technologies, U.S.). Bacterial culture strain ATCC 27853 was grown to the mid-log phase (OD_600_ = 0.7), washed, and resuspended in the same volume of PBSM (1X PBS, 1mM MgCl_2_). The bacterial culture was treated with sub-inhibitory concentrations of compounds (1/8^th^, 1/16^th^ and 1/25^th^ MIC) for 3 h at 37°C. After treatment, the cells were lysed in chloroform which was subsequently removed by boiling at 80°C. The persistent ATP from cell lysate was measured in a 96-well black flat-bottom plate by measuring the luminescence. Luminescence was converted to concentration from standard curve. A culture treated with CCCP (12.5, 6.25 and 4 μg/mL) and Azithromycin (12.5, 6.25 and 4 μg/mL) was included as positive and negative controls. Error bars represent the standard deviation of triplicates.

### 9. Determination of Frequency of Resistance (FoR) levofloxacin resistance in *P. aeruginosa*

Frequency of resistance studies were carried out as described previously [23–24].

### 10. Identification of mutation by whole-genome sequencing

DNA extraction, library preparation and whole-genome sequencing of *P. aeruginosa* parent strains ATCC 27853 and DA7232, and single isolates EPIR1S, EPIR9S, EPIR20L, EPIR43 and EPIR24L was performed by CD Genomics (New York, NY, US) using Illumina. Illumina sequencing reads were mapped using the published genome of *P. aeruginosa* ATCC 27853 (GenBank accession number CP011857), or the published genome of *P. aeruginosa* PAO1 (parent strain of DA7232) as reference genomes with the BWA-MEM tool from the Galaxy web platform (https://usegalaxy.org/) [24]. Variations in the genomes between resistant strains and parent strains were identified using the LoFreq tool from the same platform.

### 11. Time-kill studies

Time-kill studies were carried out as described previously [23–24] with the following changes. When indicated, moxifloxacin was added to the prepared bacterial suspensions at one time the MIC (2 μg/mL). TXA09155 was added to bacterial suspensions at 1/8^th^, 1/12.5^th^, and 1/16^th^ times the MIC (6.25 μg/mL, 4 μg/mL, and 3.125 μg/mL, respectively).

### 12. Determination of PAE by measuring turbidity

Exponentially grown *P. aeruginosa* ATCC27853 was incubated with levofloxacin (1x MIC) alone, and combined with 6.25 μg/mL of TXA11114 for 1 h at 37°C [64]. Treated cultures suspensions were diluted 50-fold to eliminate any drug carry over. Prior growth resumption, untreated cell concentration was adjusted to equal to treated samples to minimize difference in inoculum size. An aliquot of 200 μL was loaded to a 96-well flat-bottom microtiter plates in duplicate sets and optical density was measured periodically 15 minutes intervals. PAE was derived from PAE= T_50_ – C_50_, where T_50_ and C_50_ are the time in hours required for the drug-treated and untreated cultures, respectively, to reach a value of OD_600nm_ corresponding to 50% of the final absorbance reached by an untreated control [65].

### 13. Ion Channel Method

The automated whole cell patch-clamp (Qpatch HT) technique is used to record depolarizing currents, hNav1.5 and hCav1.2, and repolarizing potassium currents, hERG. Cells Recombinant HEK-293 cells stably transfected with human Nav1.5 cDNA, Recombinant HEK293 cell line expressing the human Cav1.2 (L-type voltage-gated calcium channel, hCav1.2 α1C/β2a/α2δ1, and recombinant CHO-K1 cells stably transfected with human hERG cDNA are used separately in each of these assays. The cells are harvested by Accutase and maintained in Serum Free Medium at room temperature before assay. On the instrument the cells are pipetted into each well of a 48-well plate in external solution. Test concentrations Stock solution is prepared in DMSO at 300x the final assay concentrations, and stored at −20 °C until the day of assay. On the day of the assay, an aliquot of the stock solution is thawed and diluted into external solution to make final test concentrations. A final concentration of 0.33% DMSO is maintained for each concentration of the assay compounds and controls. The assay is conducted at room temperature. Recording conditions hNav1.5 Peak current Onset and steady state block of peak Nav1.5 current is measured using a pulse pattern, repeated every 5 sec, consisting of a hyperpolarizing pulse to −120 mV for a 200 ms duration, depolarization to −15 mV amplitude for a 40 ms duration, followed by step to 40mV for 200 ms and finally a 100 ms ramp (1.2 V/s) to a holding potential of −80 mV. hCav1.2 Currents are evoked following a 100 ms pulse to – 60 mV followed by a 50ms pulse to +10mV before returning to the holding potential of −90 mV. This paradigm is delivered three times once every 20 s. Cells are held at −90 mV with a 5 s pulse to −60 mV every 20 s for a total of 120 s between each set of three pulses. The cell is held at −80 mV. Then the cell is depolarized to +40 mV for 500 ms and then to −80 mV over a 100ms ramp to elicit the hERG tail current. This paradigm is delivered once every 8s. The Extracellular Solution (control) is applied first and the cell is stabilized in the solution for 5 min. Then the test compound is applied from low to high concentrations sequentially on the same cell. The cells are incubated with each test concentration for 5 min. Reference compounds Tetracaine Nifedipine, and E-4031 are tested concurrently for hNav1.5, hCav1.2 and hERG, respectively at multiple concentrations to obtain an IC50 value.

hNav1.5 The amplitude of the sodium current is calculated by measuring the difference between the peak inward current on stepping to −10mV (i.e. peak current) and the leak current. The sodium current is assessed in vehicle control conditions and then at the end of each five (5) minute compound application. Individual cell result is normalized to the vehicle control amplitude and the mean ± SEM calculated for each compound concentration. These values are then plotted and estimated IC50 curve fits calculated. hCav1.2 The maximum inward current elicited on stepping to +10 mV for 50msec from −60mV is measured. Compound interaction assessed by dividing Post current amplitude (at the end of each three (3) minute compound application) by the vehicle control amplitude and the mean calculated for each compound concentration. These values are then plotted and estimated IC50 curve fits calculated. The percent inhibition of hERG channel is calculated by comparing the peak (hERG tail) current amplitude before and after application of the compound (the current difference is normalized to vehicle control values).

## Synthetic Procedures and Characterization Data

### General Methods

All Chemicals and solvents were used as received from the vendors without prior treatment or purifications. The analysis process was thorough and comprehensive. Unless otherwise stated, thin layer chromatography (TLC) was done on 0.25 nm thick precoated silica gel 60 (HF-254, Whatman). Flash column Chromatography (FCC) was performed using Kieselgel 60 (32-63 mesh, Scientific Adsorbents). Elution for FCC usually employed a stepwise solvent polarity gradient correlated with TLC mobility. Chromatograms were observed under UV (short and long wavelength) light and/ or were visualized by heating plates that were exposed to iodine/silica mix, dipped in a basic potassium permanganate, KMnO_4_ stain solution, dipped in phosphomolybdic acid stain solution, or dipped in alcoholic Ninhydrin solution. 1H NMR and ^13^C and ^19^F NMR Spectra were recorded using Mercury Varian 300 MHz instrument (300 MHz, 75 MHz, and 282MHz respectively) using CDCl_3_, Pyridine-*d_5_*, MeOH-*d_4_*, or DMSO solution with residual CDCl_3_, Pyridine-*d_5_*, MeOH-*d_4_*, or DMSO as internal standard. Chemical shifts are quoted in (δ ppm) relative to the corresponding solvent peak and a coupling constant (*J*) are given in Hertz, multiplicity (s =singlet, d= doublet, t = triplet, q = quartet, m = multiplet). Data for ^13^C are reported as follow: chemical shift (δ ppm), multiplicity (s = singlet, d = doublet, t = triplet, q = quartet, m = multiplet) and coupling constant *J* (Hertz, Hz). Low-resolution mass spectrometry was performed on Shimadzu LC-MS system using gradient mobile phase of 0.01% formic acid solution water/CH_3_CN. Analyses were performed using Shimadzu technology LCMS 2020 system and the use of analytical column Phenomenex®, 4.6 mm Gemini-NX 3u C18 110 A 50×4.6 mm). Chromatography was performed at ambient temperature with run time = 6 min with flow rate of 0.8 mL/ min with linear gradient Water (0.1% formic acid): CH_3_CN (0.1% formic acid [100:0] to water (0.1% formic acid): CH_3_CN (0.1% formic acid) [10:90] and resolved peaks detected by SPD-20AV photodiode Array (PDA) Detector at 254, 280, and/or 215 nm and characterized by low-resolution mass spectrometry instrument (Shimadzu) with ESI ion source and positive mode ionization.

### Synthesis of Intermediates 5-10

1.1 Synthetic protocols for intermediate alcohol **5** (di-*tert*-butyl (4*R*)-2-fluoro-5-hydroxypentane-1,4-diyl)dicarbamate)

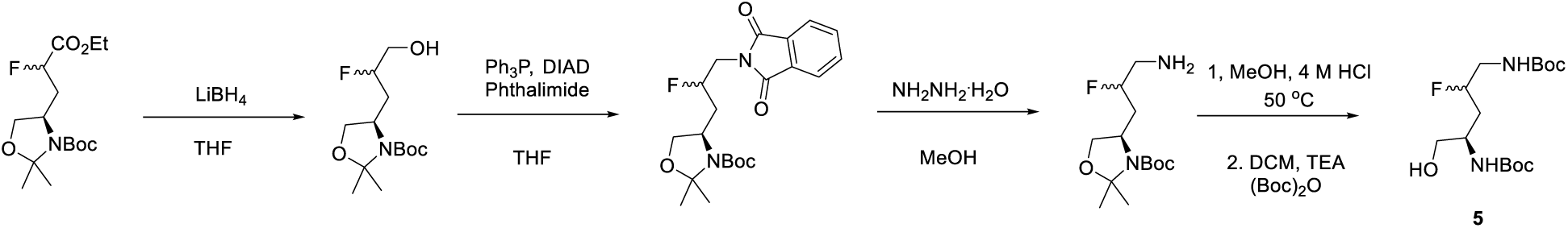

To a solution of the known ester tert-butyl *(4S)-*4-(3-ethoxy-2-fluoro-3-oxopropyl)-2,2-dimethyloxazolidine-3-carboxylate (10.8 g, 33.8 mmol) in THF (150 mL) at room temperature was added LiBH_4_ (1.62 g, 74.4 mmol) portion-wise. The reaction mixture was stirred at room temperature overnight then quenched by the slow addition of acetone in portions and the resulting mixture was concentrated to give a residue. The residue was diluted with EtOAc and washed with H_2_O, saturated NaHCO_3_, and brine. The organic solution was dried over Na_2_SO_4_, filtered, and concentrated to give the crude product. The crude product was purified on silica gel and elution with 50% EtOAc/Hexanes to afford the desired product (8.54 g, 91% yield). ^1^H NMR (300 MHz, CDCl_3_) *δ* 4.68 (dm, *J* = 54 Hz, 1H, CHF), 3.62 - 4.03 (m, 5H), 1.83 - 2.23 (m, 2H), 1.41 - 1.62 (m, 15H).

To a solution of the resulting alcohol from the previous step *tert*-butyl (4S)-4-(2-fluoro-3-hydroxypropyl)-2,2-dimethyloxazolidine-3-carboxylate (6.36 g, 22.9 mmol) in dry THF (100 mL) was added triphenylphosphine (6.61 g, 25.2 mmol), phthalimide (3.71 g, 25.2 mmol), and DIAD (5.1 mL, 25.2 mmol) at 0 °C. The reaction mixture was stirred at 0 °C and gradually warmed up to room temperature overnight. The reaction mixture was concentrated and purified on column chromatography on silica gel using gradient elution 5-20% EtOAc/hexanes to afford the desired product (7.64 g, 82% yield) as a white solid. NMR values are listed for the major isomer ^1^H NMR (300 MHz, CDCl_3_) *δ* 7.85 (m, 2H), 7.72 (m, 2H), 4.87 (dm, *J* = 50.4 Hz, 1H, CHF), 4.02-3.67 (m, 5H), 2.25-1.89 (m, 2H), 1.53 (m, 6H), 1.45 (m, 9H). ^13^C NMR (75 MHz, CDCl_3_) *δ* 168.2, 152.5, 134.8, 132.2, 123.2, 93.3, 90.1 (d*, J* = 173.2 Hz), 80.5, 67.9, 56.1, 42.2, 36.7, 28.6, 27.8, 24.8.

To a solution of the corresponding phthalimide *tert-*butyl (4*S*)-4-(3-(1,3-dioxoisoindolin-2-yl)-2-fluoropropyl)-2,2-dimethyloxazolidine-3-carboxylate (6.3 g, 15.5 mmol) in MeOH (100 mL) was added hydrazine monohydrate (2.35 mL, 31.0 mmol). The mixture was stirred at room temperature overnight. The formed precipitate was filtered off and washed with CH_2_Cl_2_. The filtrate was concentrated and triturated with CH_2_Cl_2_. The solid was removed by filtration. The filtrate was washed with saturated NaHCO_3,_ brine and dried over Na_2_SO_4,_ filtered, then concentrated and purified on silica gel column. Elution with EtOAc then 10% MeOH/CH_2_Cl_2_ with 1% NH_3_.H_2_O afforded the product (3.68 g, 86% yield) as a colorless gum. NMR values are listed for the major isomer, ^1^H NMR (300 MHz, MeOD-*d_4_*) *δ* 4.61 (dm, *J* = 50.7 Hz, 1H), 4.1 (m, 2H), 3.88 (m, 1H), 2.83 (m, 2H), 1.94 (m, 2H), 1.56 (s, 3H), 1.51 (m, 12H). ^13^C NMR (75 MHz, MeOD-d_4_) *δ* 154.0, 95.1 (d, *J* = 165 Hz), 94.5, 81.9, 69.2, 57.4, 47.2, 37.4, 28.9, 28.1, 25.0.

To a solution of the resulting fluoro-amine from the previous step *tert-*butyl (4*S*)-4-(3-amino-2-fluoropropyl)-2,2-dimethyloxazolidine-3-carboxylate (3.23 g, 11.7 mmol) in MeOH (50 mL) was added 4 N HCl solution in dioxane (11.7 mL, 46.8 mmol). The reaction mixture was stirred at 50 °C for 1 h then concentrated to give a residue. The residue was dissolved in MeOH/ CH_2_Cl_2_ (10 mL/100 mL) then added TEA (6.5 mL, 46.8 mmol), (Boc)_2_O (6.4 g, 29.3 mmol), stirred at room temperature for 3 h, then concentrated to afford a residue. The residue was diluted with EtOAc and washed with H_2_O, 10% citric acid, saturated NaHCO_3_, and brine. The organic solution was dried over Na_2_SO_4_, filtered, and concentrated to give a crude product. The crude product was purified by column chromatography on silica gel using 50%EtOAc/Hexanes to afford the desired product (2.36 g, 60% yield) as a white solid. NMR values are listed for the major isomer, ^1^H NMR (300 MHz, CDCl_3_) *δ* 5.04 (m, 2H), 4.65 (dm, *J* = 51 Hz, 1H, CHF), 3.82 (m, 1H), 3.61 (m, 2H), 3.40 (m, 1H), 3.19 (m, 1H), 2.78 (m, 1H), 1.78-1.61 (m, 2H), 1.40 (s, 18H). (75 MHz, CDCl_3_) *δ* 156.3, 91.0 (d, *J* = 168.5 Hz), 80.0, 65.6, 49.6, 44.9 (d, *J* = 20 Hz), 34.4 (d, *J* = 19.5 Hz), 28.5.

1.2 Synthetic protocols for the amine intermediate **8** (di-*tert*-butyl ((4*R*)-5-amino-2-fluoropentane-1,4-diyl)dicarbamate)

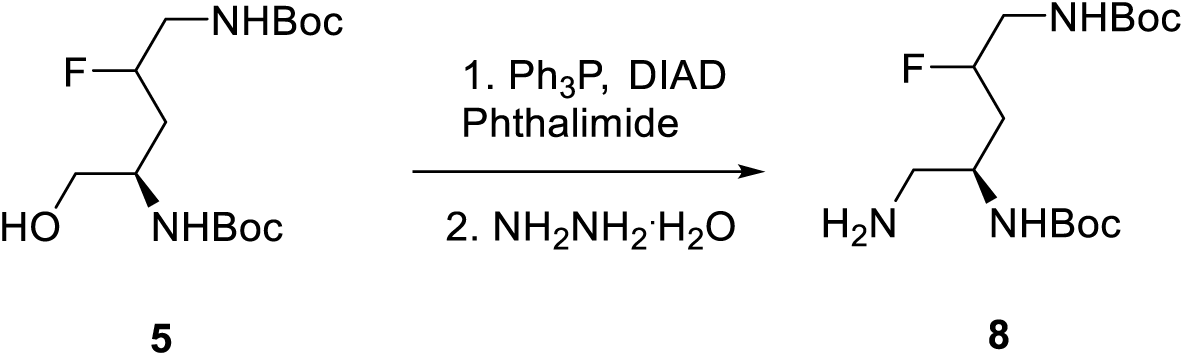

To a solution of resulting alcohol **5** *di-tert-butyl ((4R)-2-fluoro-5-hydroxypentane-1,4-diyl)dicarbamate* (2.0 g, 5.90 mmol), triphenylphosphine (1.71 g, 6.54 mmol) and phthalimide (0.96 g, 6.54 mmol) in THF (30 mL) was added DIAD (1.32 mL, 6.54 mmol) at 0 °C The reaction mixture was stirred at 0 °C then room temperature overnight. The reaction mixture was concentrated and purified on column chromatography on silica gel using 5-20% EtOAc/hexanes to give the desired product (2.54 g, 91% yield) as a white solid. ^1^H NMR (300 MHz, CDCl_3_) δ 7.83 (m, 2H), 7.70 (m, 2H), 4.82 (m, 2H), 4.70 d (m, *J* = 48 Hz, 1H, CHF), 4.20 (bs, 1H), 3.74 (m, 2H), 3.45 (m, 1H), 3.24 (m, 1H), 1.86 (m, 2H), 1.42 (s, 9H), 1.22 (s, 9H). ^1^H NMR (75 MHz, CDCl_3_) *δ* 168.5, 156.1, 155.7, 134.1, 132.2, 123.5, 90.4 (d, *J* = 169 Hz), 79.79, 79.5, 47.0, 44.7 (d*, J* = 21.2 Hz), 42.4, 35.3 (d, *J* = 20.2 Hz), 28.5, 28.2.

To a solution of the resulting phthalimide from the previous step *di-tert-*butyl ((4S)-5-(1,3-dioxoisoindolin-2-yl)-2-fluoropentane-1,4-diyl)dicarbamate (2.2 g, 4.73 mmol) in MeOH (40mL) was added hydrazine monohydrate (0.72 mL, 9.46 mmol). The mixture was stirred at room temperature overnight and the precipitate formed was filtered off and washed with CH_2_Cl_2_. The filtrate was concentrated and diluted with CH_2_Cl_2_, washed with saturated NaHCO_3,_ brine and dried over Na_2_SO_4._ The organic solution was filtered and concentrated to give a crude product. The crude product was purified on silica gel column chromatography. Elution with EtOAc then 10% MeOH/CH_2_Cl_2_ with 1% NH_3_.H_2_O afforded the product (1.30 g, 82% yield) as a colorless gum. ^1^H NMR (300 MHz, CDCl_3_) δ 4.55(dm, *J* = 54 Hz, 1h, CHF), 3.74 (bs, 1H), 3.25 (m, 2H), 2.65 (m, 2H), 1.86 - 1.66 (m, 2H), 1.44 (s, 18H).

1.3 Synthetic protocols for the alcohol intermediate **6** (di-*tert*-butyl ((4S)-3-fluoro-5-hydroxypentane-1,4-diyl)dicarbamate)

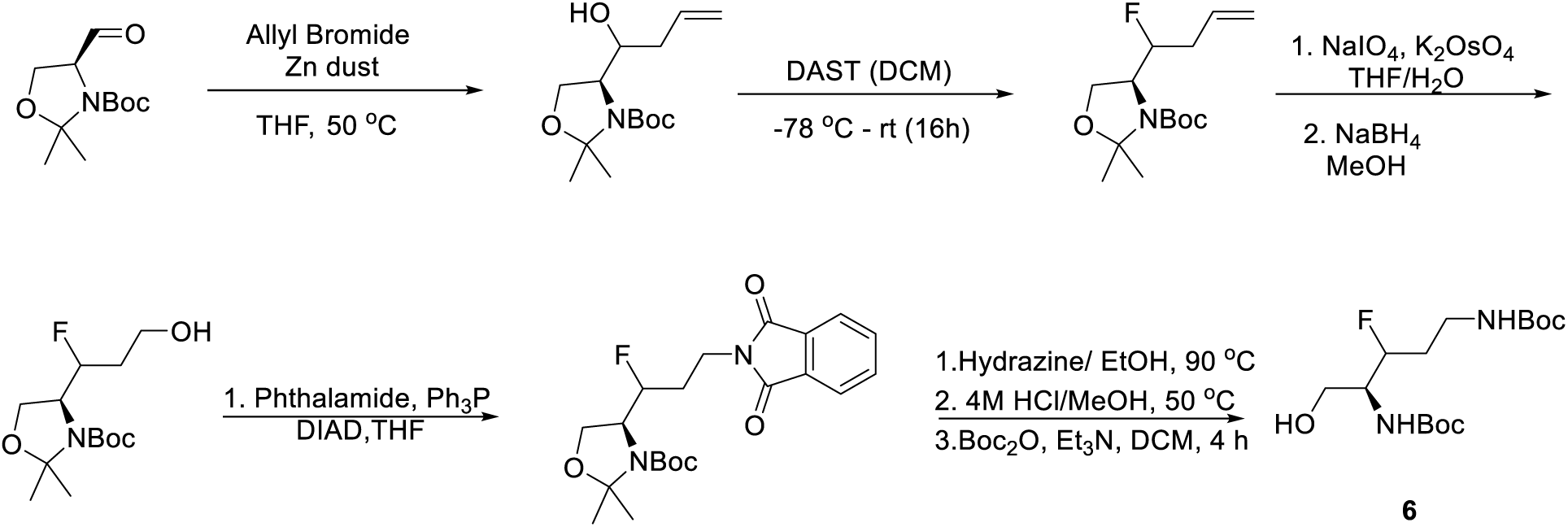

*tert*-butyl (4*S*)-4-(1-hydroxybut-3-en-1-yl)-2,2-dimethyloxazolidine-3-carboxylate:

To a solution of Garner’s Aldehyde (1.2 g, 5.2 mmol) in anhydrous THF (20 mL) was added at 0 °C Allyl bromide (540 uL, 6.2 mmol), activated Zinc dust (400 mg, 6.2 mmol) and LiCl (260 mg, 6.2 mmol). The reaction mixture was heated to 50 °C and stirred at this temperature for another 6 h. The reaction mixture was filtered over a pad of celite, and the resulting filtrate was mixed with ethanol amine (6.2 mmol). The resulting white precipitate was filtered over a short pad of silica and the resulting filtrate was concentrated under vacuum and the resulting crude was purified using ISCO flash chromatography system and the use of gradient elution of Hexanes/EtOAc (R*_f_* = 0.19 10% EtOAc/Hexane) to give 910 mg of heavy clear oil 65% yield. ^1^HNMR (300 MHz, CDCl_3_) δ 5.82 (m, 1H), 5.06 (m, 2H), 3.89 (m, 4H), 2.14 (m, 2H), 1.42 (m, 15H). ^13^CNMR (75 MHz, CDCl_3_) δ 153.8, 135.3, 117.3, 94.2, 80.8, 71.9, 64.4, 61.8, 37.9, 28.4, 26.6.

tert-butyl (4S)-4-(1-fluorobut-3-en-1-yl)-2,2-dimethyloxazolidine-3-carboxylate:

To a solution of the *tert*-butyl (4*S*)-4-(1-hydroxybut-3-en-1-yl)-2,2-dimethyloxazolidine-3-carboxylate (900 mg, 3.3 mmol) in anhydrous DCM (20.0 mL) was added at −78 C DAST (890 uL, 6.6 mL). The reaction mixture was stirred at this temperature for 1 h then gradually warmed up to rt and stirred at this temperature overnight. The reaction mixture was then treated with sat. solution of NaHCO_3_ (30 mL) And the reaction mixture was transferred into a separatory funnel.

The organic layer was washed with brine, dried over anhydrous Na_2_SO_4_. Filtered and evaporated to dryness. The crude material was purified using ISCO flash chromatography system using gradient hexanes/EtOAc elution. The target collected as a clear oil (575 mg, 64%). NMR analysis showed that the compound exists as a mixture of rotamers, the listed values represent the major rotamer. ^1^HNMR (300 MHz, CDCl_3_) δ 5.80 (m, 1H), 5.09 (m, 2H), 4.76 (dm, *J* = 48 Hz, 1H, CHF), 4.08-3.85 (m, 3H), 2.35 (m, 2H), 1.45 (m, 15H). ^13^CNMR (75 MHz, CDCl_3_) δ 153.0, 133.2, 118.4, 94.1, 91.6 (d, *J*= 173.3 Hz, CHF), 80.8, 63.2, 59.6, 37.0 (d, *J* = 26.7 Hz), 28.6, 27.0, 24.9. ^19^FNMR (282 MHz, CDCl_3_) δ −193.85 (major rotamer).

(2*S*)-2-amino-3-fluorohex-5-en-1-ol:

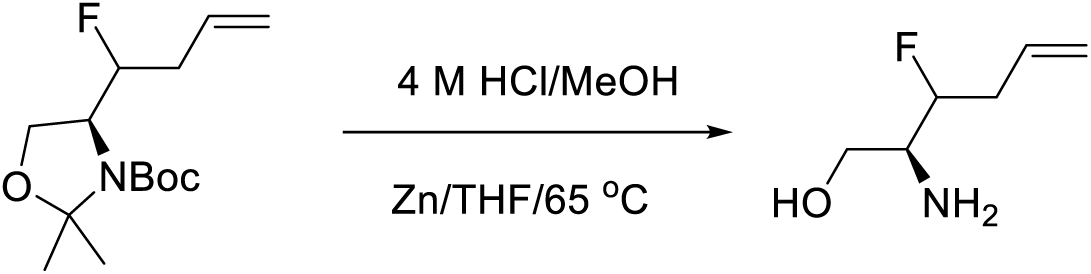

In effort to establish the diastereomeric ratio of the fluorinated derivative from the previous reaction, a (27.3 mg, 0.1 mmol) of the fluoro-alkene derivative produced in the previous step was treated with 4M HCl (500 uL), the mixture was stirred at 45 °C for 1 h, then the volatiles were removed under vacuum and the crude material was dissolved in THF (2 mL) and treated with activated zinc dust (20 mg, 0.3 mmol). The reaction mixture was heated at 65 oC for additional hour, then the reaction mixture was filtered over short pad of celite and the filtrate was evaporated to dryness, the resulting solid was analyzed as a crude mixture to establish the ratio which found almost 4:1, the listed values are for the major isomer. ^1^HNMR (300 MHz, MeOD-*d_4_*) δ 5.90 (m, 1H), 5.18 (m, 2H), 4.75 (dm, *J* = 46.8 Hz, 1H), 3.94 (m, 1H), 3.67 (m, 2H), 3.23 (m, 1H), 2.51 (m, 2H).^13^CNMR (75 MHz, MeOD*d_4_*) δ 134.0, 119.0, 94.0 (d, *J* = 172.7 Hz, CHF), 60.7, 56.4 (d, *J* = 21.0 Hz), 36.8 (d, *J* = 21.1 Hz).

tert-butyl (4S)-4-(1-fluoro-3-hydroxypropyl)-2,2-dimethyloxazolidine-3-carboxylate:

To a solution of the *tert*-butyl (4*S*)-4-(1-fluorobut-3-en-1-yl)-2,2-dimethyloxazolidine-3-carboxylate from the previous step (330 mg, 1.2 mmol) in THF (35.0 mL) was added water (15.0 mL), NaIO_4_ (775.6 mg, 3.6 mmol) and K_2_OsO_2_.2H_2_O (9.0 mg, 0.025 mmol). The reaction mixture was stirred at 0 °C and left warming gradually to rt overnight. After 16 h standing at rt, the reaction mixture was treated with saturated solution of sodium sulfite (50 ml) and stirred vigorously at rt for 2 h, then the mixture was transferred to a separatory funnel and mixed with EtOAc (50 mL), the aqueous layer was extracted with EtOAc (25 mL X 3). The combined organic layer was washed with brine (50 mL), dried over anhydrous Na_2_SO_4_, filtered, and evaporated under vacuum; the crude material was used in the next step without further purification. NMR values are listed for the crude material without purification. ^1^HNMR (300 MHz, CDCl_3_) δ 9.77 (m, 1H), 5.05 (dm, *J* = 45 Hz, 1H, CHF), 4.08-3.92 (m, 3H), 2.82 (m, 2H), 1.45 (m, 15H). ^13^CNMR (75 MHz, CDCl_3_) δ 198.6, 158.9, 94.4, 87.7 (d, *J= 172.5 Hz*, CHF), 81.2, 64.1, 59.4 (*d, J = 27.8 Hz*), 46.5 (d, *J = 21.2 Hz*), 28.5, 27.5, 24.5. The crude aldehyde from the previous step was dissolved in Methanol (20.0 mL) and treated with NaBH_4_ (133.2 mg, 3.6 mmol). The reaction mixture was then stirred at rt for 30 min before it was quenched with saturated solution of ammonium chloride. The aqueous layer was extracted with EtOAc (25 mL X 3) and the organic layer was washed with brine (25 mL), dried over anhydrous Na_2_SO_4_, filtered, and evaporated under vacuum. The resulting crude material was used in the next step without further purification. The primary alcohol was collected as clear oil (290 mg, 87% over two steps). The NMR values are listed for the major rotamer. ^1^HNMR (300 MHz, CDCl_3_) δ 4.81 (dm, J = 46.2 Hz, 1H), 4.07 (m, 2H), 3.91 (m, 2H), 3.80 (m, 2H), 1.86 (m, 2H), 1.46 (m, 15H). ^13^CNMR (75 MHz, CDCl_3_) δ 153.0, 94.1, 90.9 (d, J = 178.8 Hz, CHF), 81.1, 63.5, 59.1, 35.0 (d, *J* = 22.2 Hz), 29.8 (d, *J* = 17.7 Hz) 28.6, 27.2, 24.8. ^19^FNMR (282 MHz, CDCl_3_)

δ −194.23 (major rotamer).

*tert*-butyl (4*S*)-4-(3-(1,3-dioxoisoindolin-2-yl)-1-fluoropropyl)-2,2-dimethyloxazolidine-3-carboxylate:

To a solution of the *tert*-butyl (4*S*)-4-(1-fluoro-3-hydroxypropyl)-2,2-dimethyloxazolidine-3-carboxylate (190.0 mg, 0.68 mmol) in dry THF (10 mL) was added triphenylphosphine (359.0 mg, 1.37 mmol) and Phthalimide (201.0 mg, 1.37 mmol). The reaction mixture was cooled down to 0 C using an ice bath. After 15 min, a solution of DIAD (276.0 mg, 1.37 mmol) in anhydrous THF (5 mL) was introduced dropwise to the reaction mixture over 30 min. Upon complete addition of the DIAD solution, the reaction mixture was left to warm up to rt and stirred for two hours. The reaction progress was monitored by TLC or LCMS; when the reaction came to completion, the reaction mixture was treated with MeOH (5.0 mL), and the volatiles were removed under vacuum; the crude mixture was loaded to a pre-backed RediSep column and purified using flash chromatography on an ISCO machine using gradient elution of EtOAc/Hexane to collect the target phthalimide 225 mg, 82% yield. The NMR analysis showed that the compound exists as a mixture of rotamers, the listed values are for the major rotamer. ^1^HNMR (300 MHz, CDCl_3_) δ 7.71 (m, 2H), 7.59 (m, 2H), 4.63 (dm, *J* = 46.8 Hz, 1H),3.96-3.75 (m, 5H), 1.87 (m, 2H), 1.36 (m, 12H), 1.29 (m, 3H). ^13^CNMR (75 MHz, CDCl_3_) δ 168.0, 152.7, 134.0, 132.0, 123.3, 93.9, 90.6 (d, *J* = 173.8 Hz, CHF), 80.6, 63.2, 59.6 (d, *J* = 24.5 Hz), 34.7, 31.0 (d, *J* = 20.4 Hz) 28.3, 27.0, 24.5.

^19^FNMR (282 MHz, CDCl_3_) δ −194.11 (major rotamer).

di-*tert*-butyl ((4*S*)-3-fluoro-5-hydroxypentane-1,4-diyl)dicarbamate:

To a solution of the *tert*-butyl (4*S*)-4-(3-(1,3-dioxoisoindolin-2-yl)-1-fluoropropyl)-2,2-dimethyloxazolidine-3-carboxylate (225 mg, 0.55 mmol) in EtOH (10.0 mL) was added Hydrazine hydrate (83 uL, 1.66 mmol). The reaction mixture was heated and stirred at 90 °C for 2 h; the reaction progress was monitored by LCMS; once the Starting material was consumed entirely based on the LCMS analysis, the reaction was stopped by evaporating all volatiles under a vacuum. Then, the resulting free amine was treated with 4M HCl/MeOH (4.0 mL) to cleave the acid-labile-protecting groups. The reaction mixture was stirred at 50 °C for two hours. Then, the volatiles were evaporated entirely under a high vacuum for 2 h. The Crude mixture was then suspended in DCM (20 mL) and treated with Boc_2_O (355 mg, 1.66 mmol) and Et_3_N (460 uL, 3.36 mmol). The reaction mixture was stirred at rt for three hours. Then, the reaction mixture was diluted with 20 mL of DCM and transferred into a separatory funnel; the organic layer was washed with 1 M NaOH solution (20 mL), Saturated NH_4_Cl solution (20 mL), and brine (20 mL). The organic layer was dried over Na_2_SO_4_, filtered, and concentrated under a vacuum. The resulting crude material was loaded into a RediSep silica Column and purified by flash chromatography using an ISCO machine and a gradient elution of EtOAc/Hexane. The final compound was collected after evaporating the desired fractions as a thick, clear gummy material in a 95% yield over a three-step sequence. In some cases, the final compound solidifies on standing as white crystals. NMR analysis showed that the compound exists as a mixture of diastereomers, the listed values are for the major diastereomers. ^1^HNMR (300 MHz, CDCl_3_) δ 5.32 (m, 2H), 4.94 (m, 1H), 4.58 (dm, *J* = 45.9 Hz, 1H), 3.72 (m, 2H), 3.23 (m, 2H), 1.85 (m, 2H), 1.36 (m, 18H). ^13^CNMR (75 MHz, CDCl_3_) δ 168.4, 156.2, 156.0, 91.6 (d, *J* = 172 Hz, CHF), 79.8, 79.3, 6.7 (d, *J* = 4 Hz), 54.5 (d, *J* = 24.8Hz), 37.2, 32.1 (d, *J* = 19.7Hz), 28.4, 28.3. ^19^FNMR (282 MHz, CDCl_3_) δ −191.23 (major diastereomer). HRMS m/z calculated for C_15_H_30_FN_2_O_5_ [M+H] 337.2139 found 337.2146.

1.4 Synthetic protocols for the synthesis of amine intermediate **9** (di-*tert*-butyl ((4S)-5-amino-3-fluoropentane-1,4-diyl)dicarbamate)

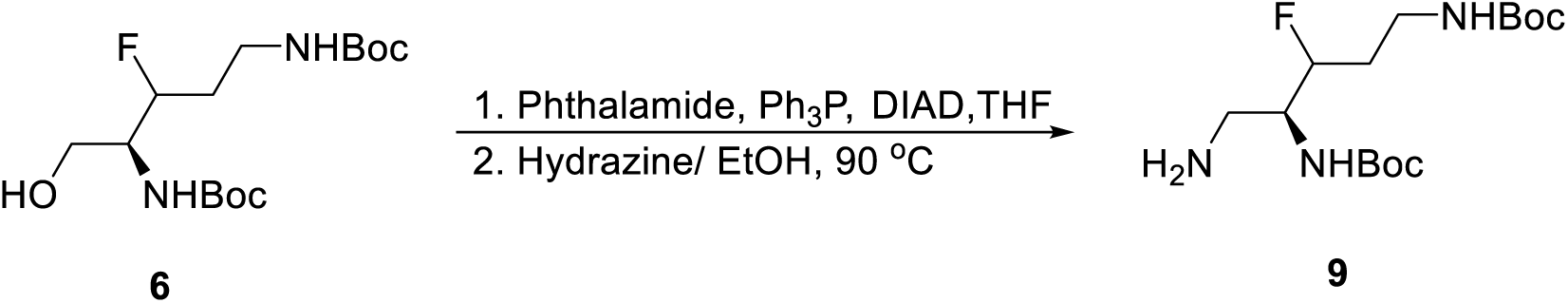

di-tert-butyl ((4S)-5-(1,3-dioxoisoindolin-2-yl)-3-fluoropentane-1,4-diyl)dicarbamate:

To a solution of the di-tert-butyl ((4S)-3-fluoro-5-hydroxypentane-1,4-diyl)dicarbamate (370 mg, 1.1 mmol) in dry THF (10.0 mL) was added triphenylphosphine (577 mg, 2.2 mmol) and Phthalimide (323 mg, 2.2 mmol). The reaction mixture was cooled down to 0 °C using an ice bath. After 15 min, a solution of DIAD (444.4 mg, 2.2 mmol) in anhydrous THF (2.0 mL) was introduced dropwise to the reaction mixture over 30 min. Upon complete addition of the DIAD solution, the reaction mixture was left to warm up to rt and stirred for another two hours. The reaction progress was monitored by TLC or LCMS; when the reaction came to completion, the reaction mixture was treated with MeOH (5.0 mL), and the volatiles were removed under vacuum; the crude mixture was dissolved in 2 mL of DCM and loaded to a pre-backed RediSep column and purified using flash chromatography on an ISCO machine using gradient elution of EtOAc/Hexane to collect the target phthalimide (114 mg, 85% yield as white solid). ^1^HNMR (300 MHz, CDCl_3_) δ 7.82 (m, 2H), 7.69 (m, 2H), 5.02 (m, 2H), 4.62 (dm, *J* = 48.5 Hz, 1H), 4.05 (m, 1H), 3.85 (m, 2H), 3.31 (m, 1H), 1.95 (m, 2H), 1.41 (m, 9H) 1.16 (m, 9H). ^13^CNMR (75 MHz, CDCl_3_) δ 168.6,, 155.7, 134.2, 132.2, 123.5, 93.4 (d, *J* = 174.8 Hz, CHF), 80.0, 79.6, 52.5 (d, *J* = 26.4 Hz), 37.5 (d, *J* = 26.4 Hz), 32.5 (d, *J* = 26.3 Hz), 28.6, 28.2.

di-*tert*-butyl ((4S)-5-amino-3-fluoropentane-1,4-diyl)dicarbamate **9**:

To a solution of the corresponding phthalimide derivative from the previous step (465 mg, 1.0 mmol) in EtOH (20 mL)) was added Hydrazine (150 uL, 3.0 mmol). The reaction mixture was stirred at 90 °C for two hours. Then, the volatiles were removed under a vacuum, and the crude material was dissolved in a minimum amount of 10% DCM/MeOH solution and loaded into a RediSep column. The material was then purified using flash chromatography using an ISCO machine and gradient elution of MeOH/DCM. The desired fractions were collected and evaporated to dryness to give a white solid with a 69% yield (over two steps). A 10 mg sample was dissolved in deuterated MeOH and analyzed by NMR spectroscopy to show mixture of diastereomers, the NMR values listed are for the major isomer. ^1^HNMR (300 MHz, MeOD-*d_4_*) δ 4.60 (dm, *J* = 46.6 Hz, 1H), 3.77 (m, 1H), 3.34 (m, 2H), 3.02 (m, 1H), 2.82 (m, 1H), 1.91 (m, 2H), 1.61 (m, 9H), 1.58 (m, 9H). ^13^CNMR (75 MHz, MeOD*d_4_*) δ 158.5, 158.3, 93.5 (d, *J* = 171.1 Hz, CHF), 80.5, 80.1, 57.1 (d, *J* = 23.6 Hz), 42.3, 37.9, 33.5 (d, *J* = 21 Hz), 29.0, 28.9. ^19^FNMR (282 MHz, MeODd_4_) δ −192.94. LRMS m/z calculated for C_15_H_31_FN_3_O_4_ [M+H] 336.2299 found 336.2308.

1.5 Synthetic protocol of the synthesis of difluoro alcohol intermediate **7** (di-*tert*-butyl (2,2-difluoro-5-hydroxypentane-1,4-diyl)(*R*)-dicarbamate).

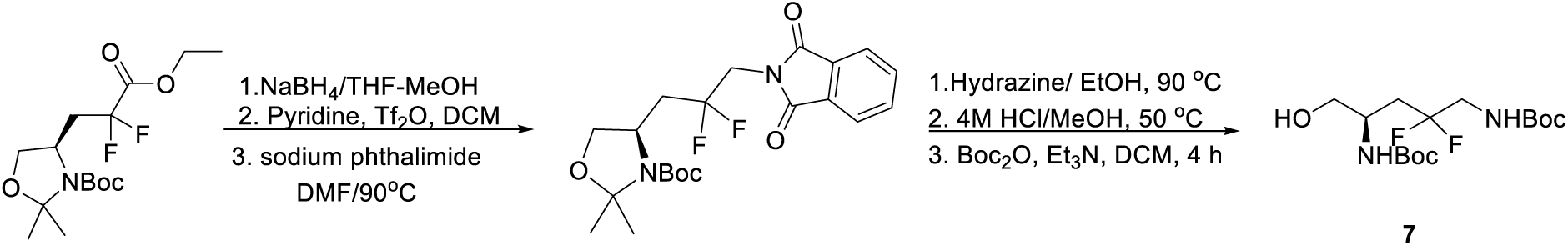

*tert*-butyl (*R*)-4-(3-(1,3-dioxoisoindolin-2-yl)-2,2-difluoropropyl)-2,2-dimethyloxazolidine-3-carboxylate:

To a solution of ester **4** *tert*-butyl (*R*)-4-(3-ethoxy-2,2-difluoro-3-oxopropyl)-2,2-dimethyloxazolidine-3-carboxylate (1012 mg, 3.0 mmol) in 5:1 solution of THF/MeOH (20 mL) was added at 0 °C NaBH_4_ (555 mg, 15 mmol). The reaction mixture was stirred at this temperature for 30 min and then warmed up gradually to rt and the stirring continued for another 3 h. Upon the complete conversion based on the tlc, the reaction mixture was treated with NH_4_Cl saturated solution at 0 °C and the reaction mixture was stirred for additional hour, the solid precipitate was filtered off and the filtrate was transferred to a separatory funnel and extracted with EtOAc (25 mL x 3), the combined organic layer was washed with brine, dried over Na_2_SO_4_, filtered, and evaporated under vacuum. The resulting crude was passed through a short silica column with EtOAc elution. The excess EtOAc was evaporated, and the crude alcohol was used in the next step without purification. 1HNMR (300 MHz, C6D6) δ 3.91 (m, 2H), 3.66 (m, 3H), 2.57-2.08 (m, 2H), 1.45 (s,3H), 1.41 (s, 3H), 1.27 (s, 9H).

To a solution of the corresponding alcohol from the previous step tert-butyl (R)-4-(2,2-difluoro-3-hydroxypropyl)-2,2-dimethyloxazolidine-3-carboxylate (500 mg, 1.67 mmol) in DCM 2.0 mL, was added pyridine (157 uL, 1.99 mmol) at 0 °C, followed by dropwise addition of Tf_2_O solution (336 uL, 1.99 mmol) in 2.0 mL of DCM. The reaction mixture was stirred at 0 °C for 30 min, warmed up gradually to rt, and left stirring at this temperature for another 45 min. The reaction mixture was mixed with 10 mL DCM and 1M HCl (10 mL), and the organic layer was then washed with water and Brine, dried over Na_2_SO_4_, filtered, and evaporated to dryness. The resulting triflate was used without further purification in the next step. To a solution of phthalimide (49 mg, 0.33 mmol) in anhydrous DMF (2 mL) was added sodium hydride, NaH (60% dispersed on mineral oil) (13.2 mg, 0.33 mmol), the reaction mixture was stirred under N_2_ gas at rt for 45 min, the formed sodium phthalimide was transferred using cannula to the triflate derivative made in the previous step (119 mg, 0.27 mmol) in DMF (2.0 mL), the reaction mixture was then heated to a 125 °C under N_2_ gas for 24 h. Upon completion and consumption of all the triflate starting material based on the LCMS monitoring, the reaction mixture was cooled down to rt and mixed with water 10 mL and EtOAc (10 mL). The reaction mixture was transferred into a separatory funnel, the aqueous layer was extracted by EtOAc (10 mL x 3). The combined organic layer was washed with 1 M HCl (10 mL) and brine (10 mL), dried over Na_2_SO_4_, filtered, and evaporated under vacuum to give the crude product, which was subjected to FCC using ISCO chromatography system and EtOAc/Hexane mobile phase. The desired phthalimide product was collected as a white solid (41 mg, 35%) R*_f_* = 0.25 (20%EtOAC/Hexane). NMR analysis showed that the compound exists as a mixture of rotamers, the listed spectroscopic values are for the major rotamer. ^1^HNMR (300 MHz, CDCl_3_) δ 7.87 (m, 2H), 7.74 (m, 2H), 4.24-3.96 (m, 3H), 3.91 (t, *J = 9.0 Hz*, 2H), 2.52-2.21 (m, 2H), 1.55 (m, 3H), 1.46 (m, 9H), 1.37 (m, 3H). ^13^CNMR (75 MHz, CDCl_3_) δ 167.6, 154.0, 152.0, 134.6, 132.0, 123.9, 121.4 (t, *J = 242 Hz, CF_2_*). 93.7, 80.7, 80.3, 68.0, 52.6, 42.6, 38.7, 28.6, 27.0, 23.3.

di-tert-butyl (2,2-difluoro-5-hydroxypentane-1,4-diyl)(R)-dicarbamate **7**:

To a solution of the *tert*-butyl (*R*)-4-(3-(1,3-dioxoisoindolin-2-yl)-2,2-difluoropropyl)-2,2-dimethyloxazolidine-3-carboxylate (240 mg, 0.54 mmol) in EtOH (10.0 mL) was added Hydrazine hydrate (70 uL, 1.13 mmol). The reaction mixture was heated and stirred at 90 C for 2 h; the reaction progress was monitored by LCMS; once the Starting material was consumed entirely based on the LCMS analysis, the reaction was stopped by evaporating all volatiles under a vacuum. Then, the resulting free amine was treated with 4M HCl/MeOH (5.0 mL) to cleave the acid-labile-protecting groups. The reaction mixture was stirred at 50 °C for three hours. Then, the volatiles were evaporated entirely under a high vacuum for 2 h. The Crude mixture was then suspended in DCM (20 mL) and treated with Boc_2_O (480 mg, 2.2 mmol) and Et_3_N (610 uL, 4.4 mmol). The reaction mixture was stirred at rt for 4h. Then, the reaction mixture was diluted with 20 mL of DCM and transferred into a separatory funnel; the organic layer was washed with 1 M NaOH solution (20 mL), Saturated NH_4_Cl solution (20 mL), and brine (20 mL). The organic layer was dried over Na_2_SO_4_, filtered, and concentrated under a vacuum. The resulting crude material was loaded into a RediSep silica Column and purified by flash chromatography using an ISCO machine and a gradient elution of EtOAc/Hexane. The final compound was collected after evaporating the desired fractions as a thick, clear gummy material in a 50% yield over a three-step sequence. In some cases, the final compound solidifies on standing as white crystals. ^1^HNMR (300 MHz, CDCl_3_) δ 5.22 (m, 1H), 5.13 (m, 1H), 3.91 (m, 1H), 3.62 (m, 2H), 3.48 (m, 2H), 2.95 (m, 1H), 2.12 (m, 2H), 1.40 (m, 18H). ^13^CNMR (75 MHz, CDCl_3_) δ 156.1, 122.7 (t, *J = 241.7 Hz, CF_2_*), 80.5, 80.1, 65.1, 48.1, 45.2 (t, *J = 31.3 Hz*, CH_2_-CF_2_), 35.2 (t, *J = 23.18 Hz*, CH_2_-CF_2_), 28.54, 28.47. ^19^FNMR (282 MHz, CDCl_3_) d −101.85 HRMS m/z calculated for C_15_H_29_F_2_N_2_O_5_ [M+H] 355.2045 found 355.2053.

1.6 Synthetic protocols for the synthesis of the amin-difluoro derivative **10** (di-tert-butyl (5-amino-2,2-difluoropentane-1,4-diyl)(*R*)-dicarbamate)

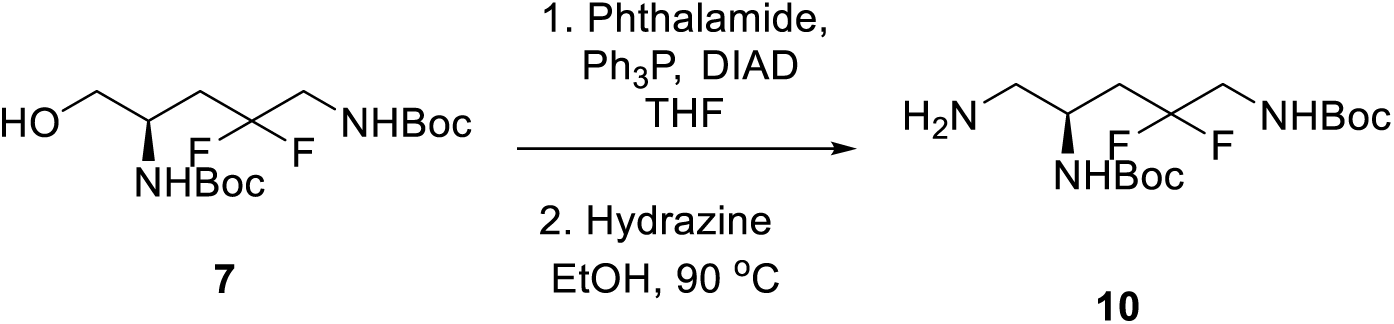

tert-butyl (R)-4-(3-(1,3-dioxoisoindolin-2-yl)-2,2-difluoropropyl)-2,2-dimethyloxazolidine-3-carboxylate:

To a solution of the di-tert-butyl (2,2-difluoro-5-hydroxypentane-1,4-diyl)(R)-dicarbamate (100 mg, 0.28 mmol) in dry THF (5.0 mL) was added triphenylphosphine (148 mg, 0.565 mmol) and Phthalimide (84 mg, 0.565 mmol). The reaction mixture was cooled down to 0 °C using an ice bath. After 15 min, a solution of DIAD (114.1 mg, 0.565 mmol) in anhydrous THF (2.0 mL) was introduced dropwise to the reaction mixture over 30 min. Upon complete addition of the DIAD solution, the reaction mixture was left to warm up to rt and stirred for another two hours. The reaction progress was monitored by TLC or LCMS; when the reaction came to completion, the reaction mixture was treated with MeOH (5.0 mL), and the volatiles were removed under vacuum; the crude mixture was dissolved in 2 mL of DCM and loaded to a pre-backed RediSep column and purified using flash chromatography on an ISCO machine using gradient elution of EtOAc/Hexane to collect the target phthalimide (114 mg, 85% yield as white solid). ^1^HNMR (300 MHz, CDCl_3_) δ 5.7.81 (m, 2H), 7.68 (m, 2H), 5.15 (m, 1H), 5.06 (m, 1H), 4.24 (m,1H), 3.79 (m, 2H), 3.53 (m, 2H), 2.13 (m, 2H), 1.39 (m, 9H), 1.23 (m, 9H). ^13^CNMR (75 MHz, CDCl_3_) δ 168.5, 156.6, 155.6, 134.2, 132.3, 123.4, 122.4 (t, *J = 240 Hz, CF_2_*), 80.5, 79.8, 45.7, 45.3 (t, *J = 30 Hz*, CH_2_-CF_2_), 42.2, 36.7 (t, *J = 24.8 Hz*, CH_2_-CF_2_), 28.5, 28.3.

(di-*tert*-butyl (5-amino-2,2-difluoropentane-1,4-diyl)(*R*)-dicarbamate) **10**:

To a solution of the di-*tert*-butyl (5-(1,3-dioxoisoindolin-2-yl)-2,2-difluoropentane-1,4-diyl) (*R*)-dicarbamate (100 mg, 0.2 mmol) in EtOH (10 mL)) was added Hydrazine (30 uL, 0.6 mmol). The reaction mixture was stirred at 90 °C for two hours. Then, the volatiles were removed under a vacuum, and the crude material was dissolved in a minimum amount of 10% DCM/MeOH solution and loaded into a RediSep column. The material was then purified using flash chromatography using an ISCO machine and gradient elution of MeOH/DCM. The desired fractions were collected and evaporated to dryness to give a white solid with a 95% yield. A 10 mg sample was dissolved in deuterated MeOH and analyzed by NMR spectroscopy. ^1^HNMR (300 MHz, MeOD-*d_4_*) δ 3.86 (m, 1H), 3.49 (m, 2H), 2.66 (m, 2H), 2.04 (m, 2H), 1.46 (m, 18H). ^13^CNMR (75 MHz, MeOD-*d_4_*) δ, 158.4, 158.1, 124.0 (t, *J* = 241.5 Hz, CF_2_), 80.7, 80.3, 49.5, 47.0, 45.9 (t, *J* = 30.8 Hz), 37.5 (t, *J* = 24.8 Hz), 28.9, 28.8. ^19^FNMR (282 MHz, CDCl_3_) d −103.42. HRMS m/z calculated for C_15_H_30_F_2_N_3_O_4_ [M+H] 354.2204 found 354.2212.

### Synthesis of Intermediates 11

2. Synthetic protocols for the carboxylic acid **11** (6-(4-fluorophenyl)-1H-indole-2-carboxylic acid)

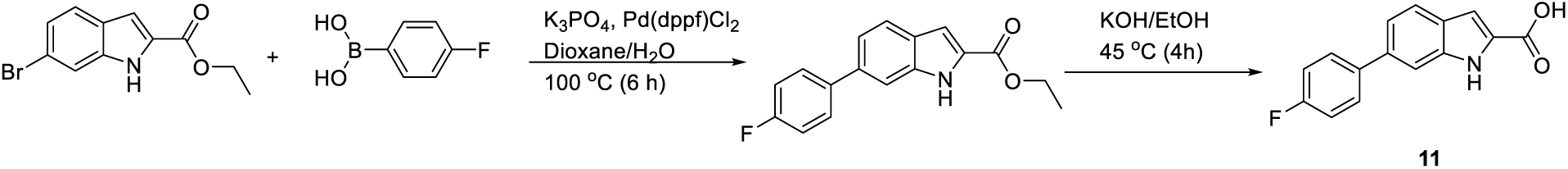

ethyl 6-(4-fluorophenyl)-1*H*-indole-2-carboxylate:

In a clean 250 mL sealed tube charged with stir bar was added Ethyl 6-bromo-1H-indole-2-carboxylate (804.3 mg, 3.0 mmol), dioxane (20 mL), (4-fluorophenylboronic acid) (840 mg, 6.0 mmol), K_3_PO_4_ (1908 mg, 9.0 mmol), and water (10.0 mL). The reaction mixture was degassed for 30 min by passing N_2_ gas through the solution. After that, Pd(dppf)Cl_2_ (122 mg, 0.15 mmol) was added and the reaction vessel was sealed and refluxed at 100 ^0^C for 6 h. Then, the reaction mixture was cooled down to rt and the volatiles were concentrated under vacuum. The resulting crude was redissolved with 50% EtOAc/Hexane. And the mixture was then filtered through short pad of silica. The organic layer was concentrated and the crude mixture was purified by flash chromatography using gradient elution of EtOAc/Hexane. The titled compound was collected as yellow solid with a 68.5% yield (*R_f_* = 0.43, 10% EtOAc/hexane).^1^HNMR (300 MHz, CDCl_3_) δ 9.21 (s, 1H), 7.72 (m, 1H), 7.56 (m, 3H), 7.34 (m, 1H), 7.24 (m, 1H), 7.13 (m, 2H), 4.43 (m, 2H), 1.43 (t, *J* = 7.11Hz, 3H). ^13^CNMR (75 MHz, CDCl_3_) δ 162.6 (d, *J = 245 Hz, Sp2-C-F*), 162.2, 138.0, 137.6, 129.1 (d, *J = 8.0 Hz*), 128.3, 128.5, 127.0, 123.1, 121.0, 115.8 (d, *J = 21 Hz*), 110.2, 108.8, 61.3, 14.6.

6-(4-fluorophenyl)-1H-indole-2-carboxylic acid **11**:

To a solution of the ethyl 6-(4-fluorophenyl)-1*H*-indole-2-carboxylate (283 mg, 1.0 mmol) in ethanol (8.0 mL) and Water (4.0 mL) was added KOH (112.2 mg, 2.0 mmol), the reaction mixture was stirred at 45 °C for 4h at which all starting material disappeared. The reaction mixture was then treated with saturated solution of NaHSO_4_ to adjust the pH of the solution to pH = 7.0. The formed precipitate was collected by vacuum filtration, washed with cold water and the collected solids were dried further inside an oven at 55 °C. The free acid was collected as white solid in 82%. ^1^HNMR (300 MHz, MeOH-*d_4_*) δ 7.67 (m, 4H), 7.33 (dd, J = 8.4, 1.5 Hz, 1H), 7.18 (m, 3H). ^13^CNMR (75 MHz, MeOH-*d_4_*) δ 165.2, 163.8 (d, *J = 243 Hz, Sp2-C-F*), 139.7, 139 (d, *J = 87 Hz*), (d, *J = 7.8 Hz*), 128.2, 123.7, 121.2, 116.6 (d, *J = 21.8 Hz*), 111.4, 109.2. ^19^FNMR (282 MHz, MeOD*d_4_*) δ -118.62.

### Synthesis and Characterization Data of TXA Compounds

3.1 Synthetic protocols for amide TXA11114: (*N-((2R*)-2,5-diamino-4-fluoropentyl)-6-(4-fluorophenyl)-1H-indole-2-carboxamide)

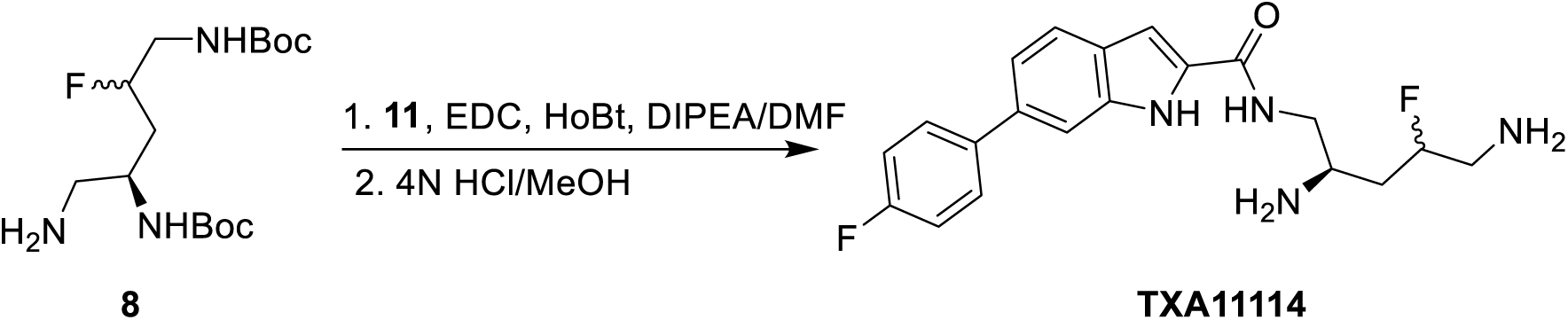

To a solution of 6-(4-fluorophenyl)-1*H*-indole-2-carboxylic acid (420 mg, 1.64 mmol) in DMF (10 mL) was added DIPEA (0.53 ml, 3.00 mmol), HOBt (120 mg, 0.89 mmol), EDC (342 mg, 1.80 mmol). The reaction mixture was stirred at room temperature then di-*tert*-butyl ((4*R*)-5-amino-2-fluoropentane-1,4-diyl)dicarbamate (500 mg, 1.50 mmol) was added and the reaction was continued to stir at room temperature overnight. The reaction mixture was diluted with EtOAc.

The combined organic layer was washed with water and brine then dried over anhydrous sodium sulfate and filtered. The filtrate was then concentrated and purified by column chromatography on silica gel using 20-30% ethyl acetate in hexanes to give the product di-*tert*-butyl ((4R)-2-fluoro-5-(6-(4-fluorophenyl)-1H-indole-2-carboxamido)pentane-1,4-diyl)dicarbamate (690 mg, 81% yield) as a white solid. ^1^H NMR (300 MHz, CDCl_3_) δ 9.21 (bs, 1H), 7.67 (d, *J* = 8.4 Hz, 1H), 7.59-7.53 (m, 3H), 7.32 (dd, *J* = 8.4, 1.6 Hz, 1H), 7.12 (t, *J* = 8.7 Hz, 1H), 6.89 (m, 1H), 5.12 (d, *J* = 8.5 Hz, 1H, NH), 4.89 (m, 1H, NH), 4.75 (dm, J = 52.2 Hz, 1H, CHF), 4.09 (m, 1H), 3.543 (m, 2H), 3.28 (m, 2H), 1.87 (m, 2H), 1.42 (s, 9H), 1.39 (s, 9H). . ^13^C NMR (300 MHz, CDCl_3_) δ 162.6 (d, *J* = 244 Hz, Sp2-CF), 162.2, 157.1, 156.3, 138.2, 137.3, 137.0, 131.4, 129.1 (d, *J* = 8.3 Hz), 127.3, 122.6, 120.8, 115.8 (d, *J* = 21.5 Hz), 110.2, 102.8, 90.9 (d, *J* = 168.8 Hz, CHF), 80.5, 80.1, 48.0, 44.8 (d, *J* = 16.7 Hz) 35.1 (d, *J* = 22.4 Hz), 28.5.

To a solution of di-*tert*-butyl ((4*R*)-2-fluoro-5-(6-(4-fluorophenyl)-1H-indole-2-carboxamido)pentane-1,4-diyl)dicarbamate (670 mg, 1.19 mmol) in MeOH (10 mL) was added HCl solution (4 M in dioxane, 1.19 mL). It was stirred at rt overnight and solvent was removed under vacuo. The residue was triturated with EtOAc and the product *N-((2R*)-2,5-diamino-4-fluoropentyl)-6-(4-fluorophenyl)-1*H*-indole-2-carboxamide: **TXA11114** was collected as a white powder (495 mg, 95% yield). ^1^H NMR (300 MHz, CD_3_OD) δ 7.73-7.67 (m, 4H), 7.37 (dd, *J* = 8.4, 1.5 Hz, 1H), 7.22-7.16 (m, 3H), 5.20 (dm, J = 51 Hz, 1H, CHF), 3.85-3.65 (m, 3H), 3.44-3.26 (m, 2H), 2.27-2.06 (m, 2H). 13CNMR (75 MHz, Me3OD) δ 165.4, 163.9 (d, *J* = 143.2 Hz, sp2 CF), 139.8, 139.2, 138.2, 132.3, 130.1 (d, *J* = 8 Hz), 128.4, 123.5, 121.3, 116.6 (d, *J* = 21.5 Hz), 111.3, 105.4, 89.2 (d, *J* = 170 Hz, Sp3CF), 50.6, 44.2 (d, *J* = 20.3 Hz),42.9, 34.3 (d, J = 19.6 Hz). MS (ESI+): 396.25 [M+H]^+^ for C_22_H_26_FN_5_O.

3.2 Synthetic protocols for the amide TXA11164: (*N*-((2*S*)-2,5-diamino-3-fluoropentyl)-6-(4-fluorophenyl)-1*H*-indole-2-carboxamide)

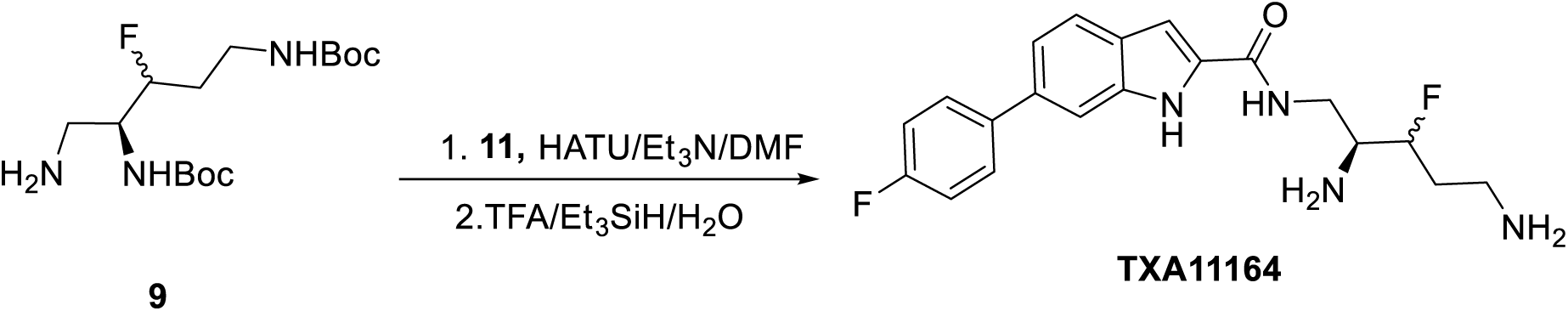

To a solution of 6-(4-fluorophenyl)-1*H*-indole-2-carboxylic acid **11** (45 mg, 0.18 mmol) in DMF (2.0 mL) was added a coupling agent HATU (137 mg, 0.0.36 mmol), and di-*tert*-butyl ((4*S*)-5-amino-3-fluoropentane-1,4-diyl) dicarbamate (60 mg, 0.18 mmol). The reaction mixture was cooled down to 0 °C then was treated with Et_3_N (100 uL, 0.74 mmol). The reaction was stirred at 0 °C for 15 min, then let warm up gradually to rt and stirred at ambient temperature for another 30 min. The reaction mixture was diluted with a saturated solution of NH_4_Cl (5.0 mL), the aqueous layer was extracted with EtOAc (10.0 mL X 3). The combined organic layer was washed with brine (5.0 mL), dried over Na_2_SO_4_, filtered, and evaporated under vacuum. The Crude material was purified using ISCO flash chromatography system and the use of gradient elution of Hexane/EtOAc. The evaporation of the desired fractions gave the di Boc protected amide. The desired amide di-tert-butyl ((4S)-3-fluoro-5-(6-(4-fluorophenyl)-1H-indole-2-carboxamido)pentane-1,4-diyl)dicarbamate was collected as white solid in 78 % yield (80 mg). ^1^HNMR (300 MHz, DMSO*d_6_*) δ 11.73 (s, 1H), 8.53 (m, 1H), 7.71 (m, 3H), 7.65 (s, 1H), 7.35 (m, 3H), 7.16 (s, 1H), 6.94 (m, 2H), 4.55 (dm, *J* = 46.6 Hz, 1H, CHF), 3.87 (m, 1H), 3.49 (m, 2H), 3.09 (m, 2H), 1.74 (m, 2H), 1.40 (m, 9H) 1.37 (m, 9H). ^13^CNMR (75 MHz, DMSO*d_6_*) δ 161.5 (d, *J* = 242.2 Hz, Sp2-CF), 161.3, 155.5, 155.4, 137.7, 137.0, 134.6, 132.4, 128.6 (d, *J* = 8.0 Hz), 126.5, 121.9, 119.2, 115.7 (d, J = 21.0 Hz), 110.0, 102.5, 91.9 (d, *J* = 170.2 Hz, CHF), 78.1, 77.5, 53.0 (d, J = 23.1 Hz), 36.3, 31.8 (d, *J* = 19.3 Hz), 28.2, 28.0.

The resulting amide from the previous step (80 mg, 0.136 mmol) was treated with 95% TFA/ 2.5% Et_3_SiH, 2.5% H_2_O solution (1.5 mL). The reaction mixture was stirred at rt for 45 min, then the volatiles were removed under vacuum to give the free amine product *N-((2S)-*2,5-diamino-3-fluoropentyl)-6-(4-fluorophenyl)-1H-indole-2-carboxamide: **TXA11164** as TFA salt in 82% yield. ^1^HNMR (300 MHz, MeOD*d_4_*) δ 7.78 (m, 4H), 7.44 (d, *J* = 8.9 Hz, 1H), 7.26 (m, 3H), 5.18 (dm, *J* = 48.3 HZ, CHF), 4.01-3.78 (m, 3H), 3.39 (m, 2H), 2.36 (m, 2H). ^13^CNMR (75 MHz, MeOD*d_4_*) δ 165.3, 163.8 (d, *J* = 243.2 Hz, Sp2-CF), 139.7 (d, *J* = 3.2 Hz), 138.2, 132.2, 130.0 (d, *J* = 7.6 Hz), 128.4, 123.4, 121.3, 116.5 (d, *J* = 21.0 Hz), 111.3, 105.5, 91.7 (d, *J* = 173.0, CHF), 55.4 (d, *J* = 20.1 Hz), 38.5, 37.6, 29.9 (d, *J* = 20.7 Hz). ^19^FNMR (282 MHz, MeODd_4_) δ −118.83, −197.73. HRMS m/z calculated for C_20_H_23_F_2_N_4_O [M+H] 373.1840 found 373.1835.

3.3 Synthetic protocols for amide TXA12027 (*R*)-*N*-(2,5-diamino-4,4-difluoropentyl)-6-(4-fluorophenyl)-1*H*-indole-2-carboxamide:

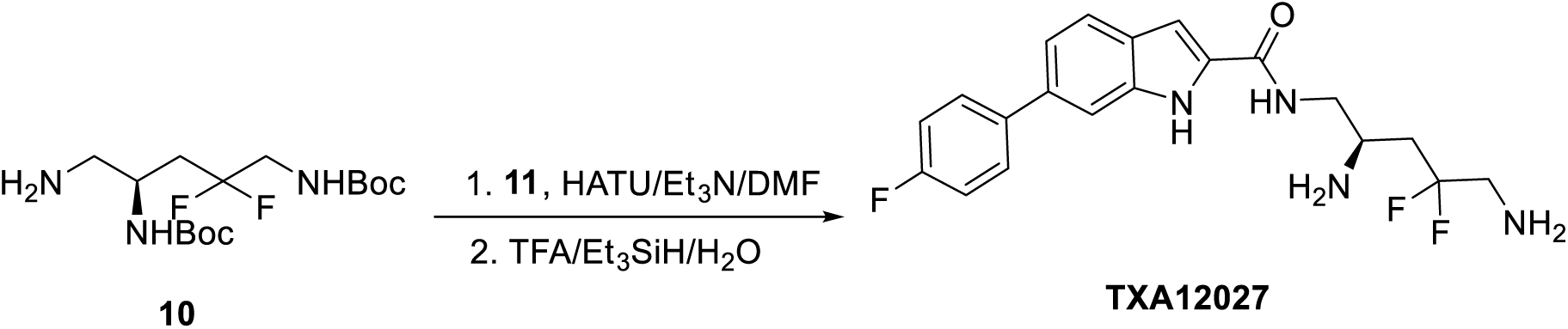

To a solution of 6-(4-fluorophenyl)-1*H*-indole-2-carboxylic acid **11** (15 mg, 0.06 mmol) in DMF (1.0 mL) was added a coupling agent HATU (22.5 mg, 0.06 mmol), and di-*tert*-butyl (5-amino-2,2-difluoropentane-1,4-diyl) (*R*)-dicarbamate (19 mg, 0.05 mmol). The reaction mixture was cooled down to 0 °C then was treated with Et_3_N (15 uL, 0.11 mmol). The reaction was stirred at 0 °C for 15 min, then let warm up gradually to rt and stirred at ambient temperature for another 30 min. The reaction mixture was diluted with a saturated solution of NH_4_Cl (2.0 mL), the aqueous layer was extracted with EtOAc (5.0 mL X 3). The combined organic layer was washed with brine (2.0 mL), dried over Na_2_SO_4_, filtered, and evaporated under vacuum. The Crude material was purified using ISCO flash chromatography system and the use of gradient elution of Hexane/EtOAc. The evaporation of the desired fractions gave the di Boc protected amide. The desired amide di-*tert*-butyl (2,2-difluoro-5-(6-(4-fluorophenyl)-1H-indole-2-carboxamido)pentane-1,4-diyl)(*R*)-dicarbamate was collected as white solid in 85 % yield (25 mg). ^1^HNMR (300 MHz, CDCl_3_) δ 9.87 (m, 1H), 7.52 (m, 5H), 7.29 (d, *J* = 8.3 Hz, 1H), 7.08 (t, *J* = 8.7, 2H), 6.95 (m, 1H), 5.66 (m, 1H), 5.06 (m, 1H), 4.18 (m, 1H), 3.54 (m, 4H), 2.12 (m, 2H), 1.40 (m, 9H) 1.35 (m, 9H). ^13^CNMR (75 MHz, CDCl_3_) δ 162.6, 162.5 (d, *J* = 244.5 Hz, Sp2-CF), 156.9, 156.1, 138.2 (d, *J* = 3.2 Hz), 137.2 (d, *J* = 9.9 Hz), 131.3, 129.0 (d, *J* = 7.9 Hz), 127.2, 122.5, 121.6 (t, *J* = 240.0 Hz), 120.7, 115.8 (d, *J* = 18.5 Hz), 110.4, 103.2, 80.7, 80.4, 46.6, 45.4, 45.3, 36.3 (t, *J* = 23.3 Hz), 28.5, 28.4.

The resulting amide from the previous step (25 mg, 0.045 mmol) was treated with 95% TFA/ 2.5% Et_3_SiH, 2.5% H_2_O solution (0.5 mL). The reaction mixture was stirred at rt for 45 min, then the volatiles were removed under vacuum to give the free amine product *(R)-N-*(2,5-diamino-4,4-difluoropentyl)-6-(4-fluorophenyl)-1H-indole-2-carboxamide: **TXA12027** as TFA salt. ^1^HNMR (300 MHz, MeOD*d_4_*) δ 7.67 (m, 4H), 7.33 (dd, *J* = 1.3, 8.3 Hz, 1H), 7.28 (s, 1H), 7.16 (t, *J* = 8.8, 2H), 3.96 (m, 1H), 3.76 (m, 4H), 2.64 (m, 2H). ^13^CNMR (75 MHz, MeOD*d_4_*) δ 165.5, 163.8 (d, *J* = 243 Hz, Sp2-CF), 139.7 (d, *J* = 3.2 Hz), 139.2, 138.2, 132.2, 130.0 (d, *J* = 7.7 Hz), 128.4, 122.5, 123.5, 122.0 (t, *J* = 243.5 Hz), 116.5 (d, *J* = 20.7 Hz), 111.3, 105.7, 48.2, 44.7 (t, *J* = 24.2 Hz), 43.2, 36.5 (t, *J* = 25.9 Hz). ^19^FNMR (282 MHz, MeODd_4_) δ −91.09, −77.42. HRMS m/z calculated for C_20_H_22_F_3_N_4_O [M+H] 391.1746 found 391.1747.

---

**Supplemental Table 1.**
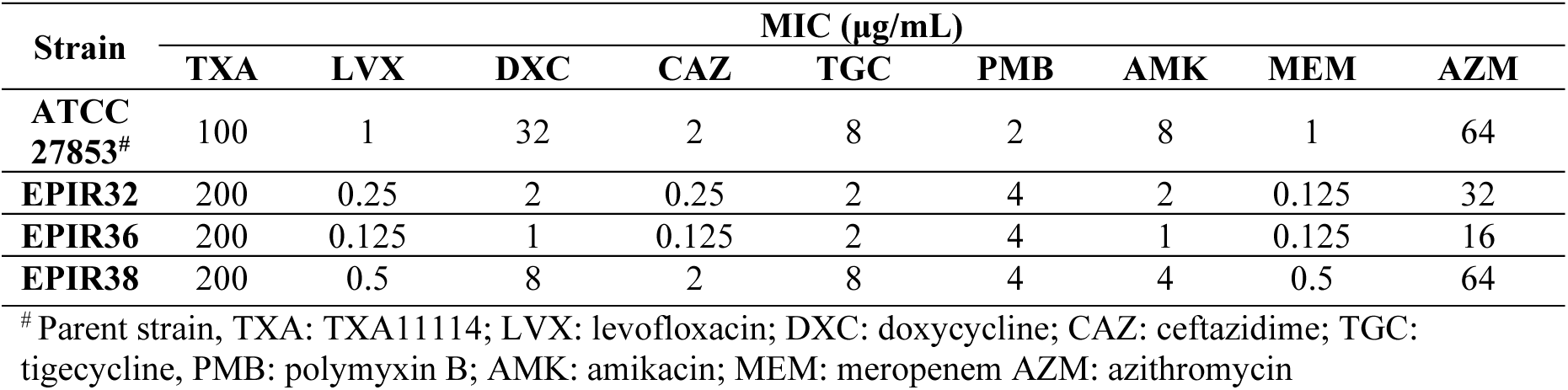
Susceptibility of *P. aeruginosa* mutants resistant to TXA11114 combination to various antimicrobials.

